# Naturally acquired antibody kinetics against *Plasmodium vivax* antigens in people from a low malaria transmission region in western Thailand

**DOI:** 10.1101/2021.10.21.464164

**Authors:** Zoe Shih-Jung Liu, Jetsumon Sattabongkot, Michael White, Sadudee Chotirat, Chalermpon Kumpitak, Eizo Takashima, Matthias Harbers, Wai-Hong Tham, Julie Healer, Chetan E Chitnis, Takafumi Tsuboi, Ivo Mueller, Rhea J Longley

## Abstract

*Plasmodium vivax* is the dominant *Plasmodium* spp. causing the disease malaria in low-transmission regions outside of Africa. These regions often feature high proportions of asymptomatic patients with sub-microscopic parasitaemia and relapses. Naturally acquired antibody responses are induced after *Plasmodium* infection, providing partial protection against high parasitaemia and clinical episodes. However, previous work has failed to address the presence and maintenance of such antibody responses to *P. vivax* particularly in low-transmission regions. We followed 34 patients in western Thailand after symptomatic *P. vivax* infections to monitor antibody kinetics over 9 months, during which no recurrent infections occurred. We assessed total IgG, IgG subclass and IgM levels to up to 52 *P. vivax* proteins every 2-4 weeks using a multiplexed Luminex^®^ assay, and identified protein-specific variation in antibody longevity. Generally, an increase in antibody level was observed within 1-week post symptomatic infection, followed by an exponential decay of different rates. We observed mostly IgG1 dominance and IgG3 sub-dominance in this population. IgM responses followed similar kinetic patterns to IgG, with some proteins unexpectedly inducing long-lived IgM responses. We also monitored antibody responses against 27 IgG-immunogenic antigens in 30 asymptomatic individuals from a similar region. Our results demonstrate that most antigens induced robust and long-lived total IgG responses following asymptomatic infections in the absence of (detected) boosting infections. Our work provides new insights into the development and maintenance of naturally acquired immunity to *P. vivax* and will guide the potential use of serology to indicate immune status and/or identify populations at risk.

## Introduction

Malaria is an infectious disease caused by parasites under the genus of *Plasmodium* and is transmitted via female *Anopheles* mosquitoes. *P. falciparum* and *P. vivax* are the two most common *Plasmodium* parasites that infect humans, causing significant morbidity and mortality worldwide. In particular, *P. vivax* is the most widely distributed *Plasmodium* species outside sub-Saharan Africa, with 2.5 billion people at risk [1, 2]. *P. vivax* is highly prevalent in southeast Asia, accounting for more than 35% of global *P. vivax* cases [3]. The capacity of *P. vivax* to cause negative impacts on human health and, in some cases, death, is significant, especially for children under 5 years of age [4, 5].

*P. vivax* has several unique features in its life cycle which make this species of *Plasmodium* a major challenge for elimination. The parasites are injected into the blood stream by a female *Anopheles* mosquito and quickly travel to the liver for maturation [6]. An unknown proportion of the liver-stage parasites remain in the hepatocytes and enter dormancy for months up to years until they are reactivated (by currently unknown signals) and re-initiating the blood-stage symptoms [7]. These arrested liver-stage parasites are known as hypnozoites, and the delayed blood-stage infections they cause are manifesting relapses of the disease [6, 7]. Relapses can contribute up to 80% of all blood-stage infections [8, 9], but no current diagnostic tools can detect individuals who have hypnozoites in their livers but no current blood-stage parasites [10]. Primaquine and Tafenoquine are available to clear hypnozoites [11], but their use is contraindicated in individuals with G6PD deficiency [12-14]. Additionally, a high proportion of low-density asymptomatic *P. vivax* infections have been reported in multiple southeast Asian regions [15-17]. These low-density asymptomatic individuals are often missed during screening and do not present to medical facilities, further contributing to ongoing transmission. New tools specifically targeting *P. vivax* are needed, such as vaccines and improved surveillance tools. Both these approaches would benefit from increased understanding of naturally acquired immune responses to the parasite.

Naturally acquired immunity is elicited upon natural exposure to pathogens and can provide significant insights into the immune status of populations [18]. In the case of infectious diseases, naturally acquired immunity is often correlated with partial protection against clinical symptoms or associated with past levels of exposure to the pathogen [19, 20]. In the context of *P. vivax* malaria, studies of naturally acquired immunity have provided novel insights such as the identification of targets of immunity for development of vaccines and sero-diagnostics [21]. Whilst the acquisition of naturally acquired total IgG responses against *P. vivax* has been relatively well studied, there have been only limited studies assessing longevity of these responses. Additionally, there is a lack of findings on the responses of IgG subclasses in low-transmission regions compared to those with higher transmission [22, 23]. It has generally been assumed that *Plasmodium*-specific antibody responses are short-lived in the absence of new boosting infections, as has been described for *P. falciparum* [19]. However, multiple studies have now reported the presence of long-lived *P. vivax*-specific antibody responses, even in the absence of ongoing exposure [24-27]. Most of these studies have focused on only a limited number of *P. vivax* antigens and thus have not assessed factors contributing to antibody longevity, with the exception of our work demonstrating that *P. vivax*-specific IgG responses were protein-specific and longer-lived in individuals from higher transmission regions [24].

To further our understandings of naturally acquired immunity and to explore its potential translational implications, we studied the acquisition and maintenance of antibody responses against a panel of 52 *P. vivax* antigens following both symptomatic and asymptomatic *P. vivax* infections in low-transmission regions in Thailand [28, 29]. Factors potentially influencing the persistence of antibody responses were also investigated. These included the presence of signal peptide (SP) [30, 31], Plasmodium exported element (PEXEL) motifs [32, 33], transmembrane domains (TMDs) [34], glycosylphosphatidylinositol (GPI) anchors [35, 36], predicted domains [37-39] and low complexity regions (LCRs) [40-42].

## Results

### Naturally acquired total IgG kinetics are antigen-specific and dominated by subclass IgG1 following symptomatic *P. vivax* infections

We first determined the levels of antigen-specific total IgG antibody in Thai symptomatic patients (n=34) at the time of *P. vivax* infection and throughout 9 months of follow-up in comparison to antibody measurements in three large malaria-naïve control panels (n=274). Antibody responses to the 52 *P. vivax* antigens were measured at baseline (week 0), weeks 1 and 2, then every 2 weeks for 6 months, followed by every month until 9 months after infection (total of 17 measurements). These individuals had no recurrent *Plasmodium* infections during the 9-month follow-up (as determined by *P. vivax*-specific PCR). An overview of the study design and the antibody measurements performed is shown in Figure 1 and Table S1. The 52 *P. vivax* antigens included both pre-erythrocytic and blood-stage expressed proteins, and were selected as they are known to be immunogenic in *P. vivax*-endemic regions, can act as markers of exposure to infection, or are of interest for vaccine development (see Methods and Table S2 for further details).

**Figure 1.**
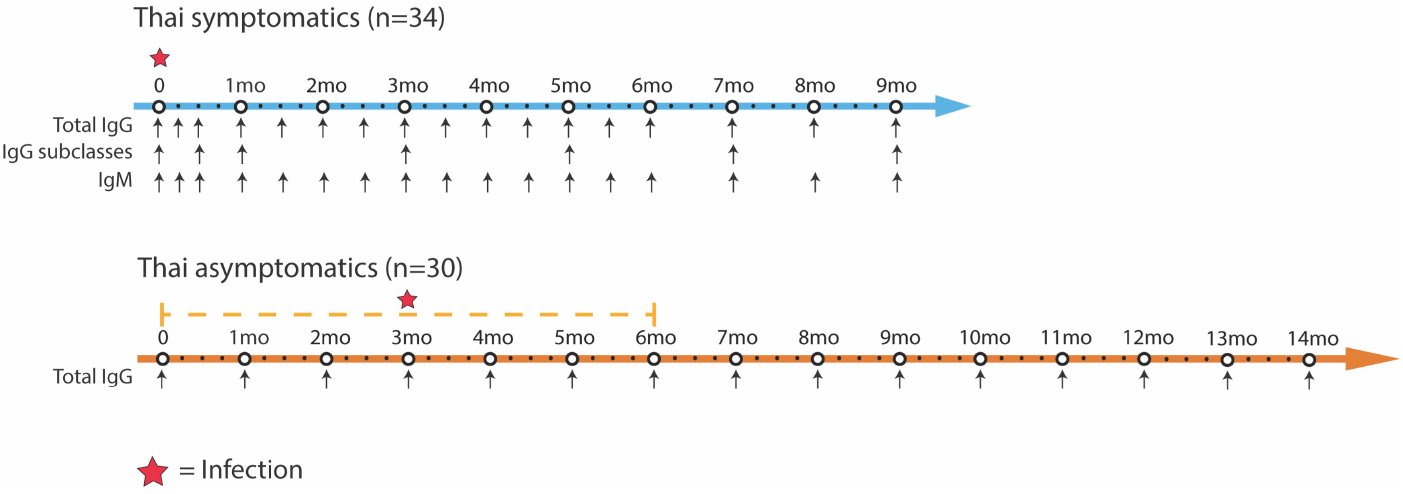
Overview of study design and antibody measurements for Thai symptomatic and asymptomatic longitudinal cohorts. The Thai symptomatic patients were all recruited at the time of clinical *P. vivax* infection then followed for 9 months. The Thai asymptomatic individuals were selected from an existing yearlong observational cohort [29]. Individuals were selected who had an asymptomatic *P. vivax* infection within the first 6 months of the study as determined by PCR during surveillance. Total IgG antibody measurements were made at the time of *P. vivax* infection and then at all available following time-points.

Total IgG antibody responses against all 52 antigens were adjusted by subtracting the negative control baseline consisting of malaria-free individuals from three separate sources (Figure 2) (the total IgG response for each individual antigen prior to adjustment is shown in Figure S1). We observed that IgG antibody responses to most *P. vivax* antigens followed a general pattern consisting of an initial peak in total IgG within two weeks post-infection and a subsequent decay of varying rates towards the baseline (Figure 2). Subsequently, we classified 27 antigens with median total IgG of more than 1 SD above the median negative control baseline at 1-week post-infection as “IgG-immunogenic” for extended analyses (Figure S1). All 27 IgG-immunogenic proteins had responses detectable above the baseline at the 9-month final visit, indicating the seropositivity against these *P. vivax* proteins is long-lived. Total IgG responses were also assessed at an individual person level, with variation evident between study participants (Figure S2). These observations could be due to differing effects of age, past exposure and history of infections with other pathogens. However, the sample size was not designed for detecting individual-level differences in IgG acquisition and maintenance.

**Figure 2.**
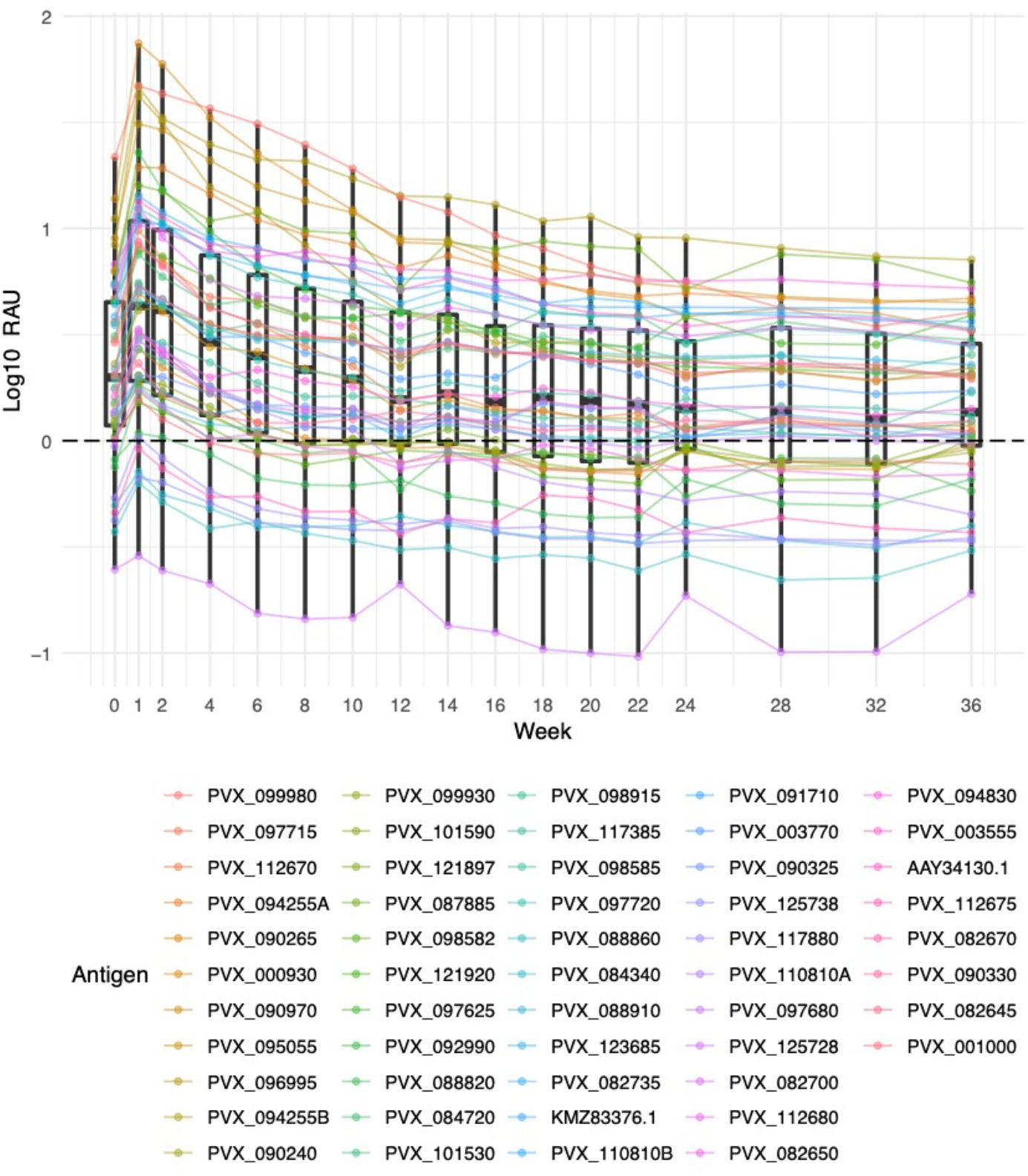
Adjusted total IgG kinetic profiles against 52 *P. vivax* antigens in 34 Thai patients with symptomatic *P. vivax* infections. Antigen-specific IgG antibody responses were measured for 9 months following a symptomatic *P. vivax* infection among all patients from western Thailand. The antibody responses were converted to arbitrary Relative Antibody Units (RAU) from Median Fluorescent Intensity (MFI) and adjusted against the median negative control baseline by subtraction. Negative control panels consist of malaria-free individuals from 3 sources: Australian Red Cross (n = 100), Thai Red Cross (n = 72) and Volunteer Blood Donor Registry (n = 102). Dots show the median of the 34 patients for each protein, box plots show the median and range of all proteins combined.

As the magnitude of the IgG response cannot be directly compared between different *P. vivax* antigens using our assay (due to differing concentrations of antigen coupled to beads), we defined immunogenicity in comparison to the hyperimmune PNG pool that was included as a positive control on each plate, as per our previous work [43]. We calculated the proportion of Thai participants, at week 1 post-infection, reaching 1, 5, 10, 25 and 50% of the antibody response in the PNG pool (Table S1). The top 5 most immunogenic proteins were PVX_000930 (sexual stage antigen s16), PVX_096995 (tryptophan-rich antigen, Pv-fam-a), PVX_097715 (hypothetical protein), PVX_112670 (unspecified protein), and PVX_094255 (reticulocyte binding protein 2b, RBP2b) (two constructs tested), with >20 of the 34 individuals reaching 50% of the antibody response in the PNG pool.

To determine the predominance and longevity of IgG subclass responses, IgG1, 2, 3 and 4 responses to 26 IgG-immunogenic antigens were measured at the time of *P. vivax* infection, week 2 post-infection and then every 2-8 weeks throughout the study follow-up (PVX_123685 was excluded from the panel by mistake) (Figure 3). Similar to total IgG, as antigen-specific baseline signals varied significantly between antigens, data were adjusted with the negative control baseline by subtraction. We observed that after adjustment IgG1 and IgG3 responses were the predominant IgG subclasses detected following symptomatic *P. vivax* infections in this western Thai population (Figure 3), with no or minimal detection of IgG2 and IgG4. IgG1 and IgG3 responses closely followed the same pattern as total IgG: an early peak followed by a decay of different rates. Of the two detectable subclasses, IgG1 was the dominant response and the IgG1 magnitude most strongly mirrored the total IgG response over time. In contrast, we found that hyper-immune adults from PNG (used as the positive control pool) had similar levels of IgG1 and IgG3 to multiple *P. vivax* proteins, and that the level of IgG3 was higher compared to that of the Thai symptomatic patients to most proteins (Figure S3).

**Figure 3.**
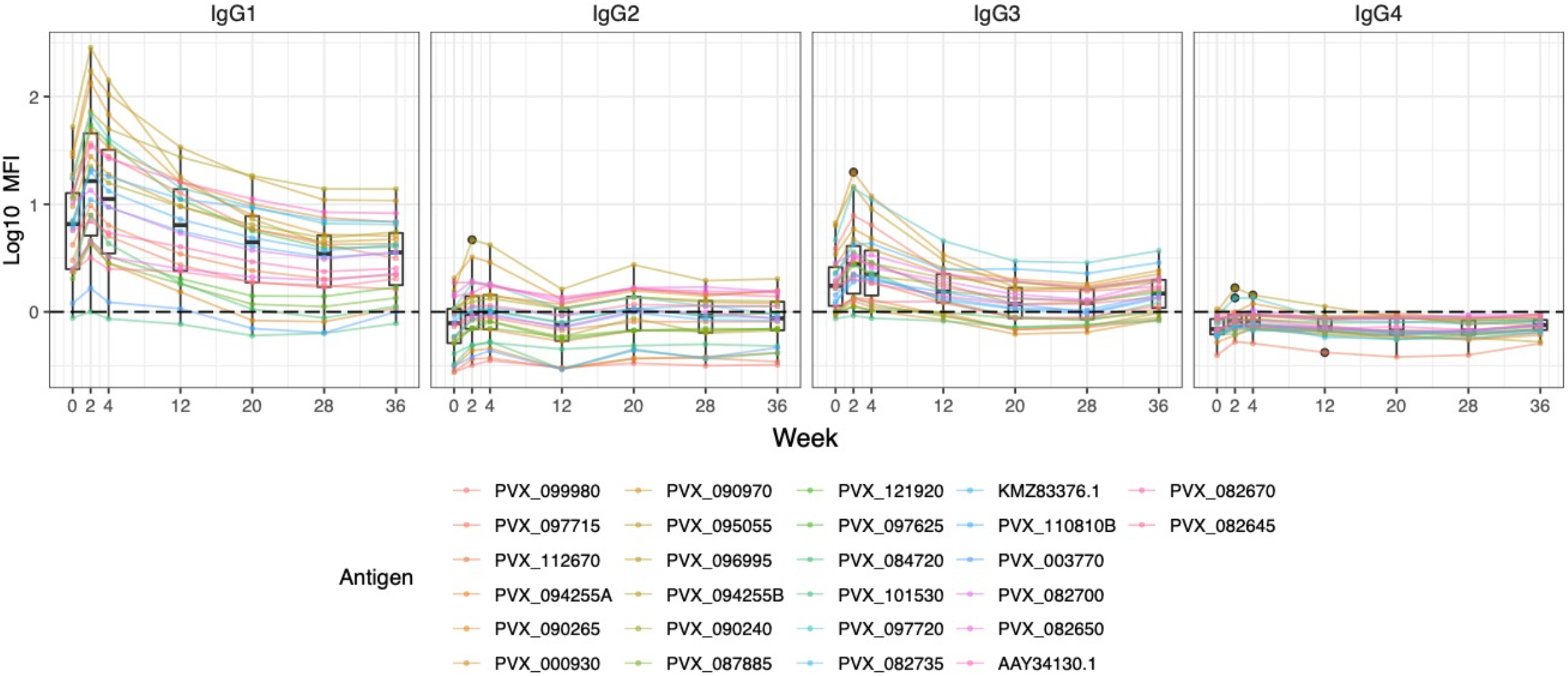
Adjusted IgG subclass kinetics in 34 Thai patients following symptomatic *P. vivax* infections. The levels of 4 IgG subclass responses against 26 total-IgG-immunogenic *P. vivax* antigens (PVX_123685 was excluded) were measured every 2-8 weeks for 9 months. Data were adjusted by subtracting the median of pooled negative control panels (n=274). Dots show the median of the 34 patients for each protein, box plots show the median and range of the population of protein

### Naturally acquired antigen-specific IgM responses are longer-lived than total IgG following symptomatic *P. vivax* infections

Whilst IgM responses are traditionally thought to be transient following infections, more recent evidence has suggested they can be long-lived or follow a similar kinetic to IgG, at least in the context of malaria [44, 45], though this is likely dependent on the antigen assessed and transmission setting [46]. We first assessed IgM immunogenicity at 1-week post-infection in the symptomatic cohort and classified 15 antigens as IgM-positive (more than 2 SD above baseline), including both IgG-immunogenic (n=8) and non-IgG-immunogenic (n=7) antigens (Figure S4). For the 15 IgM-positive antigens we assessed the kinetics over 9 months and compared the longevity to that of IgG (Figure 4), both adjusted against corresponding baseline signals of the negative control panels. Unadjusted kinetic profiles of IgM responses for each antigen are shown in Figure S5.

**Figure 4.**
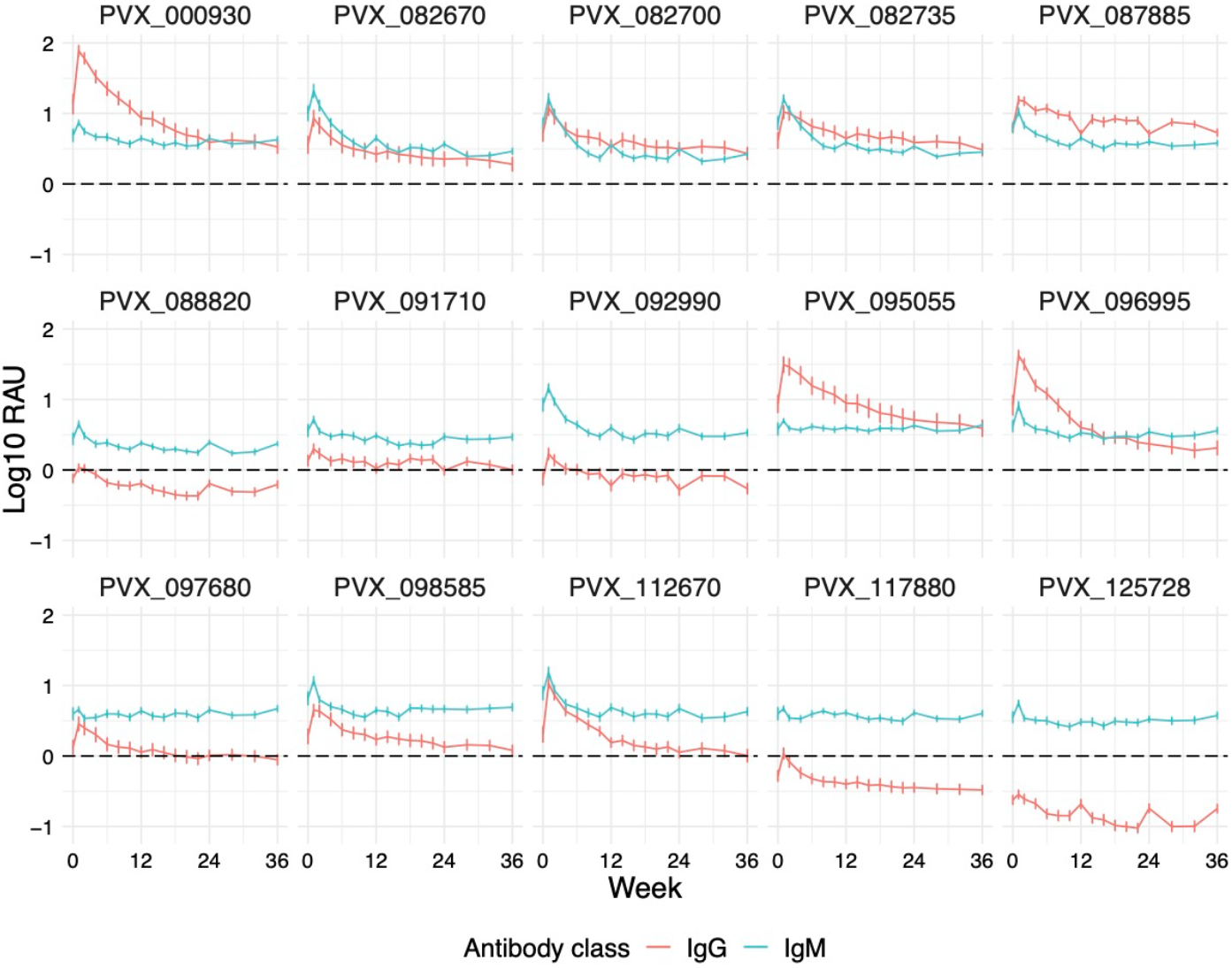
Comparison of adjusted total IgG and IgM against 15 *P. vivax* antigens following symptomatic infections. 15 antigens with IgM positivity at 1-week post-infection (> 2 SD above baseline median) were selected and their IgM responses over a 9-month period were measured and compared to total IgG responses plotted on the same scale. Data were adjusted by subtracting the median of pooled negative control panels (n=274). Data are expressed as the median ± 95% CI.

All antigens exhibited IgM kinetic patterns similar to total IgG, consisting of an initial peak at week 1 post-infection and gradual decay over the 9 months of study follow-up (Figure 4). However, the peak at week 1 post infection was generally of lesser magnitude than IgG, and thus the decay for IgG was steeper, with IgM reaching its minimal level more quickly. IgM signals against all 15 antigens were maintained above the median of negative control panels for more than 6 months, and the magnitude of the response was relatively similar for all proteins. In comparison with total IgG, variations in responses between volunteers were also less prominent in IgM (Figure 4). The relationship between IgG and IgM responses did differ between proteins, with 8 antigens exhibiting significantly higher IgM than total IgG responses throughout the study (PVX_088820, PVX_091710, PVX_092990, PVX_097680, PVX_098585, PVX_112670, PVX_117880 and PVX_125728), 4 antigens exhibiting stronger total IgG than IgM responses (PVX_000930, PVX_087885, PVX_095055 and PVX_096995) and 3 antigens where total IgG and IgM were comparable (PVX_082670, PVX_082700 and PVX_082735). The level of background IgM antibodies did vary within the malaria-naïve negative control samples tested (Figure S4), so whilst the data in Figure 4 has been adjusted against the median of the negative controls, this may not adequately account for the background cross-reactivity common for IgM and should be considered when interpreting these results. We thus focused our subsequent analyses on total IgG and IgG subclass results.

### Antigens with PEXEL motifs and no transmembrane domains are associated with longer-lived total IgG and IgG1 responses, respectively

To explore the potential sources that could account for the differences in maintenance of IgG and IgG subclass responses against *P. vivax* antigens, we examined the impact of several antigenic features that could be associated with long-lived antibody responses. These included the presence or absence of predicted signal peptides, transmembrane domains, GPI anchors, PEXEL motifs, predicted domains and the number of low complexity regions (Table S2). The majority of the 52 *P. vivax* antigens were predicted to contain signal peptides (79%), no transmembrane domains (58%), no GPI anchor (92%) and no PEXEL motif (92%). Half of the antigens were predicted to have domains. On average, these antigens had one predicted domain which was most commonly an epidermal growth factor (EGF) or Tryptophan-Threonine-rich *Plasmodium* antigen (Pfam) domain at the C-terminus. All antigens were predicted to contain LCRs (range=1-17, mean=5.98; median=4), except for PVX_098915, which had none.

In order to have a quantitative measure of antibody longevity for this analysis, we used mathematical modelling to estimate the decay rate (half-life) of antigen-specific antibodies per antigen beyond the 9-months study in the Thai symptomatic cohort (n = 34). The model of antibody kinetics was based on previous work [47, 48], with improvements as detailed in the methods. The model assumes that the antigenic stimulus from the *Plasmodium* infection results in the proliferation and differentiation of B cells into both short- and long-lived antibody secreting cells (ASCs) (or plasma cells) that secrete Ig molecules. The short- and long-lived ASCs decay at different rates resulting in the bi-phasic decay in antibody levels over time, such as observed in this study with total IgG (Figure 2). The model estimates the half-life of secreted antibody molecules and both short- and long-lived ASCs, and takes into consideration the antigen-specific background signals measured in malaria-naïve negative control panels (n=274) (Figure S6).

To assess if the presence of these antigenic features was linked to the estimated decay rate of antibodies in this cohort, we performed non-parametric Wilcoxon tests between antigens with and without the antigenic features. As shown in Figure 5A, *P. vivax* antigens with PEXEL motifs were associated with longer estimated half-life of total IgG molecules (medians = 25.72 days versus 18.20 days, p = 0.038). Additionally, antigens with no predicted TMD were associated with an estimated half-life of IgG1-producing long-lived ASCs more than three times longer than proteins with TMD domains (medians = 1,305.71 days versus 407.12 days, p = 0.037) (Figure 5B). No significant associations were observed for other antigenic features and estimated decay rates. Similarly, no significant associations were found after preforming linear regression between the number of predicted domains or LCRs and estimated half-lives generated from modelling (data not shown).

**Figure 5.**
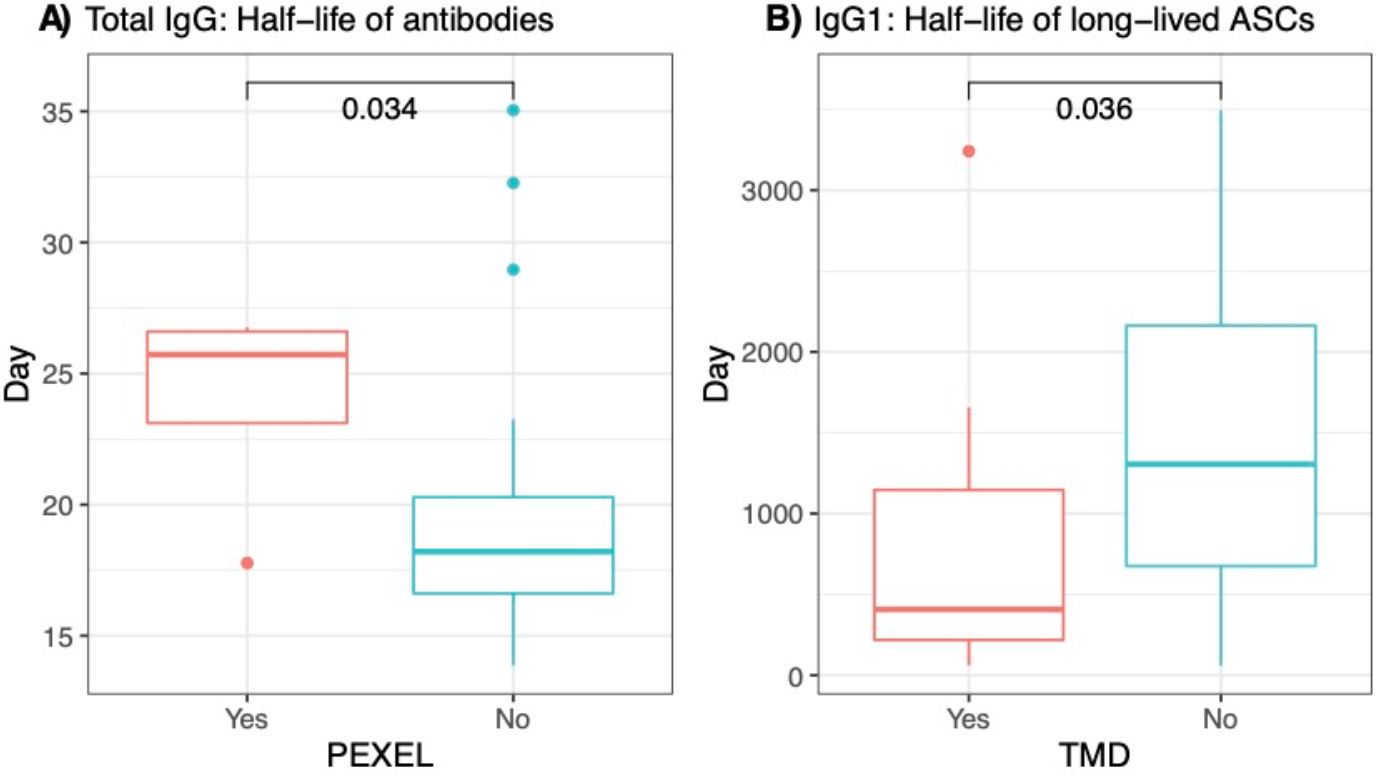
The association between presence of PEXEL motifs and estimated half-life of total IgG antibodies, and the presence of TMD and the estimated half-life of IgG1-producing long-lived ASCs in the Thai symptomatic population. The kinetics of total IgG antibody against 52 IgG-immunogenic *P. vivax* antigens were determined following symptomatic *P. vivax* infections in Thai individuals in the absence of recurrent infections for 9 months. Mathematical modelling was then used to generate the estimated half-life of total IgG and IgG1-producing long-lived ASCs. The presence of PEXEL and TMDs was predicted for each *P. vivax* antigen using the online servers: PlasmoDB (https://plasmodb.org/plasmo/app) and Transmembrane Helices Hidden Markov Model (TMHMM) Server version 2.0 (http://www.cbs.dtu.dk/services/TMHMM/). Wilcoxon tests (non-parametric) were performed to compare between antigens with (Yes) and without (No) PEXEL or TMD. p value less than 0.05 was considered significant.

### Antigen-specific total IgG responses are longer-lived following asymptomatic compared to symptomatic *P. vivax* infections

Total IgG, IgG subclass and IgM measurements were made in Thai individuals following symptomatic *P. vivax* infections. As these individuals were symptomatic, they have not yet gained sufficient immunity to clinical disease. We were therefore interested to determine whether total IgG kinetics differed in individuals from a similar region of western Thailand who had already acquired immunity to symptomatic disease. 30 individuals were selected from an existing cohort [29] who had an asymptomatic *P. vivax* infection with data available from 14 visits in total (each visit one-month apart). Each individual had data from at least 10 visits post infection, and were PCR-negative for *Plasmodium* at all of these subsequent visits. Total IgG responses were measured against the 27 *P. vivax* antigens found to be immunogenic (total IgG) in the symptomatic cohort. Kinetic profiles against these 27 antigens are shown in Figure 6 and the antigen-specific profiles in Figure S7 (with direct comparison between the symptomatic and asymptomatic cohorts).

**Figure 6.**
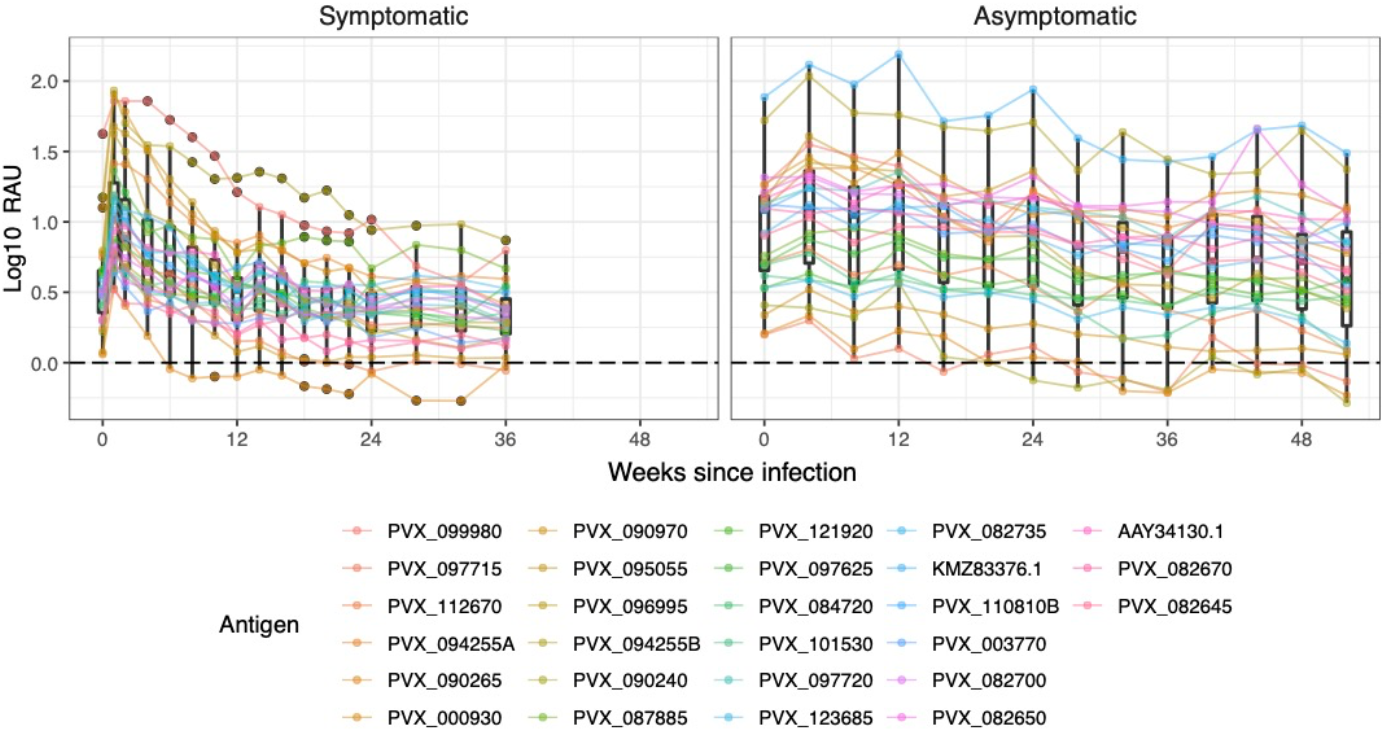
Comparison of adjusted total IgG kinetics against 27 *P. vivax* antigens following symptomatic or asymptomatic *P. vivax* infections. The kinetics of total IgG antibody against 27 IgG-immunogenic *P. vivax* antigens were determined following both asymptomatic (for a year) and symptomatic (for 9 months) *P. vivax* infections in Thai individuals in the absence of recurrent infections. Antibody data after asymptomatic infection were aligned to the time of infection, and thus sample size diminishes at later time points. Data were adjusted by subtracting the median of pooled negative control panels (n=274). Dots show the median of the 34 patients for each protein, box plots show the median and range of the population of proteins.

21 of 27 *P. vivax* antigens were found to be IgG-immunogenic (more than 1 SD above baseline at the time of infection, w0) following an asymptomatic *P. vivax* infection, all of which showed a long-lived total IgG profile (detectable IgG above baseline at 10-months post-infection). In comparison 25 proteins in the symptomatic cohort had detectable IgG at the final visit (9 months post-infection) above the median negative control baseline, with a greater drop in magnitude than observed in the asymptomatic cohort. Individual variation in responses to antigens was larger than observed in the symptomatic patients, indicated by the longer error bars in Figure S7. This is likely related to greater differences in levels of past exposure in these individuals, compared to most symptomatic patients having 0 or 1 past infections (self-reported history). For *P. vivax* antigens that were non-immunogenic in the asymptomatic *P. vivax* infected individuals (n = 6), 3 antigens (PVX_097715, PVX_090970, PVX_090265) showed high seropositivity in the symptomatic Thai cohort in which at least half (n ≥ 17) of the participants reached 50% of the antibody response in the PNG pool as shown in Table S1. The lack of response in the asymptomatic individuals could be due to the reduced parasite load; the median parasite density in the asymptomatic individuals (median = 1.0×10^2^) was 2.5×10^3^ times lower than the symptomatic (median = 2.5×10^5^) (Table 1). There was no clear peak in response at the first time-point post-infection (Figure S7), as there was for the symptomatic cohort, which could relate to the timing (monthly sampling rather than 1-2 weekly) or because the onset of infection is less well-defined in the asymptomatic cohort.

**Table 1.**
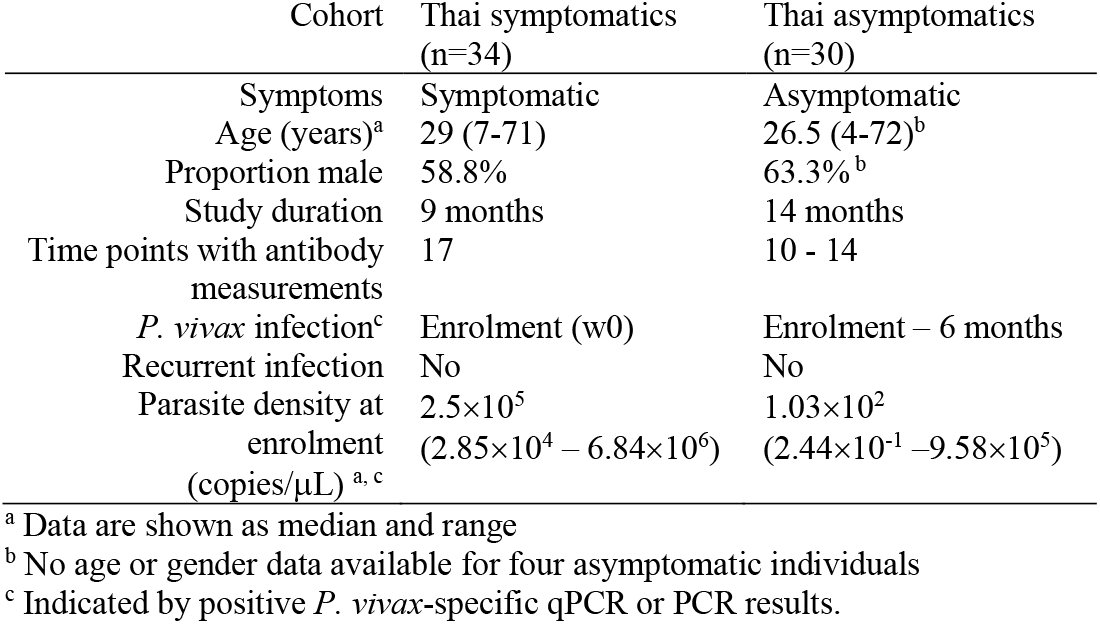
Demographic characteristics of the two study populations.

## Discussion

Antibody responses against a large number of *P. vivax* proteins are acquired following *P. vivax* infections, however little is known about how long these antibody responses last in different settings and what factors might influence the longevity. In the present study, we used human samples from a low transmission region in Thailand to develop a model of the dynamics of naturally acquired antibody responses against 52 *P. vivax* antigens, including well-studied vaccine candidates and novel targets, following both symptomatic and asymptomatic *P. vivax* infections. We show that *P. vivax* antigens are able to induce total IgG responses of varying longevities, dependent on the *P. vivax* antigen target, with a general pattern of bi-phasic decay. Of the 52 *P. vivax* antigens, 27 were highly immunogenic at the peak of the antibody response 1-week post clinical *P. vivax* infection, with all of these having detectable IgG levels above baseline at 9-months post-infection despite a decline in the magnitude. We observed that for most antigens IgG1 was the dominant subclass, and that the decay of IgG1 mirrored that of total IgG. Of interest, we also detected long-lived antigen-specific IgM responses to 15 *P. vivax* proteins, with a similar kinetic pattern after symptomatic infections. When comparing antibody kinetics following symptomatic or asymptomatic *P. vivax* infections, we show that responses following asymptomatic infections are maintained at a higher magnitude for longer, in line with our previous observations that antibody longevity increases in individuals with higher levels of past exposure [24].

To investigate the source of variation in IgG antibody longevity against different *P. vivax* antigens, we first improved upon an existing mathematical model of antibody kinetics to quantify an estimate of the half-life of secreted antibody molecules and both short- and long-lived ASCs against the 27 immunogenic proteins for total IgG, IgG1 and IgG3. We then used these quantified antibody longevity variables to assess the impact of the presence of several major protein characteristics that could influence the immune response, such as SPs, TMDs, GPI anchors and PEXEL motifs. Only two significant associations were found: 1) *P. vivax* antigens with PEXEL motifs were associated with longer estimated half-lives of total IgG molecules and 2) *P. vivax* antigens with no predicted TMD were associated with longer estimated half-lives of IgG1-producing long-lived ASCs. PEXEL motifs are responsible for trafficking and secreting of several *Plasmodium* surface antigens [49], thus proteins that contain PEXEL motifs could be capable of inducing robust immune responses, accounting for the association with longer estimated half-lives of total IgG molecules. However, there was no association between the presence of PEXEL motifs and longevity of short- or long-lived ASCs, and the difference in longevity of the IgG molecules was small (i.e. 25.72 days versus 18.20 days). We had hypothesized that the presence of TMDs would suggest the *P. vivax* antigens were membrane-bound, and thus highly accessible to the immune system. However, we found that antigens with no TMD had longer half-lives (> 3-fold) for IgG1-secreting long-lived ASCs. There was no significant relationship between the presence of TMD and total IgG or IgG3-secreting long-lived ASCs. The impact of these protein features warrants further examination, for example through the assessment of other factors that could influence antibody longevity such as regions of structural disorder [50], the presence of predicted B cell epitopes, the sequence polymorphism of the antigens, and how these interact or could confound the current findings of associations with PEXEL and TMDs. Interestingly, the only other study we could identify that has aimed to assess potential protein features and their association with antibody longevity in the context of *Plasmodium* infections, hypothesised that the tandem repeated epitope sequences in the *P. falciparum* protein MSP2 could account for the short antibody longevity observed to this protein [51] compared to the other antigens they assessed. Tandem repeats are common in regions of protein disorder, and regions of protein disorder are enriched in exported proteins [50], making this a key feature for future assessment.

Interestingly, whilst we expected the 52 *P. vivax* antigens to be mostly immunogenic based on our prior studies [24, 52], we found that only 27 induced IgG and 15 induced IgM seropositive responses at week 1 following infection. Potential reasons are that we have used purified antigens compared to our large-scale analysis of more than 300 crude *P. vivax* antigens [24], and that our sample size is much smaller than our analysis of 829 individuals from malaria-endemic regions using identical purified antigens [52]. We also set a relatively stringent cut-off of the median plus two times standard deviation for IgM, to account for the variability observed in the malaria-naïve negative controls. Other studies have also demonstrated immunogenicity against a number of our *P. vivax* antigens identified as non-immunogenic in the current study, supporting the notion that the lack of immune response identified is related to the small sample size, genetic variations between individuals and lack of past exposure in the symptomatic Thai patients. For example, whilst the *P. vivax* antigen GAMA was categorised as non-immunogenic in our study, Changrob et al. reported high seroprevalence of more than 55% even a year after recovery from acute infection in southern Thailand [53]. We classed 6 tryptophan-rich antigens as non-immunogenic, among which 2 (PVX_112675 and PVX_092990) were found to be highly immunogenic and capable of inducing long-lived antibody responses that lasted for at least 5 years post-infection in individuals from Korea, Myanmar and China [54]. Various reasons could account for such differences, in addition to those previously mentioned, including differences in study design (possibly boosting infections), protein construct sequence, detection methodology, past history of infection and region-specific *P. vivax* transmission levels.

Our data indicates that cytophilic antibodies (IgG1 and IgG3) are the dominant IgG subclass response induced by our large panel of *P. vivax* antigens, at least following symptomatic *P. vivax* infections in this low-transmission region. This is consistent with previous research highlighting IgG1 and IgG3 as the key subclasses following malaria, regardless of the *Plasmodium* species or geographical location [43, 45, 55-57]. It has previously been proposed that *P. falciparum* infection initially induces non-cytophilic followed by cytophilic antibodies [58], but this was not what we observed following *P. vivax* infections in individuals with symptomatic infections in western Thailand (who presumably have had limited past exposure, supported by their self-reported malaria history). Whether or not this observation is species-specific remains unclear. In comparison to previously published studies mostly utilising cross-sectional cohorts, our longitudinal data and relatively frequent sampling allowed us to demonstrate that the dominance of IgG subclasses remains constant over time following *P. vivax* infection, at least in the absence of any boosting infections. We did not observe any evidence of subclass switch, which is proposed to occur with an increase in exposure and age [56], in this low-transmission region. However, in our positive control pool consisting of hyperimmune individuals from PNG, we did observe the presence of both IgG1 and IgG3 against most *P. vivax* antigens, compared to IgG1 alone for our Thai volunteers, suggesting that with increasing exposure and in higher transmission levels this switch occurs. Interestingly, the subclass switch was hypothesised to contribute to the longevity of total IgG upon exposure to malaria antigens [59, 60], but was not observed in our study where the study site is of relatively low transmission.

IgM antibodies have been identified as a short-lived response due to the general notion that IgM is rapidly replaced by IgG after a short period of time exposed to a pathogen [61]. In the context of *P. vivax* infections, Park and colleagues reported *P. vivax* MSP1-specific IgM responses lasting less than 3 months [62]. However, our findings indicate that IgM responses can be long-lived (for at least 9 months), consistent with more recently published reports [63-65]. Whilst the underlying mechanism for induction and maintenance of long-lived IgM is unclear, IgM-producing long-lived plasma cells have been shown to reside in the spleen in a mouse model [64]. These cells undergo somatic hypermutation upon antigenic stimulation, independent of germinal centre and T cell help, suggesting an affinity maturation process that is unique and distinct from that of IgG [64, 66]. Our study is the first to address *P. vivax*-specific IgM kinetics in detail and to demonstrate that IgM can be maintained for at least 9 months in the absence of boosting infections. The direct comparison between total IgG and IgM kinetics also provides unique insights into the antigen-specific difference in the temporal dynamics between the two antibody responses. However, caution is warranted when interpreting these results as evidence in other disease models has shown that IgM can potentially be cross-reactive to highly IgG-immunogenic epitopes [67, 68]. Confounding factors that could potentially influence the longevity of *P. vivax*-specific IgM responses should also be taken into consideration, such as the presence of other pathogens displaying structurally similar antigens between participants in this cohort.

We also measured IgG antibodies following a PCR-positive but asymptomatic *P. vivax* infection in a similar region of western Thailand as a comparison to the symptomatic cohort in the nearby region. Defining antibody responses in asymptomatic individuals is challenging given that case detection and recruitment become more dependent on molecular surveillance tools that are less accessible than microscopy in field-settings. Yet it is important to assess longevity of naturally acquired immune responses following asymptomatic infections given the high proportion of *P. vivax* infections that are asymptomatic (i.e. [69, 70]). We were able to utilise samples previously collected as part of an observational longitudinal cohort study, in which most infections were asymptomatic [29], to address this question. We have previously shown that total IgG responses against a panel of 11 *P. vivax* antigens are induced following asymptomatic *P. vivax* infections [71]. Our current results confirm these findings, against a larger panel of *P. vivax* antigens, and we were able to further extend these results to show that these responses can be maintained for an extended period of time following clearance of the infection (> 10 months). Whilst there have been previous studies assessing antibody longevity following *P. vivax* infections [27, 53, 72, 73], ours is the first to do so to a large panel of *P. vivax* antigens with much more frequent sampling after an infection. This enables us to establish an antibody kinetic profile of higher resolution and with high certainty of no recurrent boosting infections during follow-up. In addition, the malaria-naïve control panels in our study are exceptionally larger than most previously published work, allowing more accurate prediction of seroprevalence in the target population.

Our study has some limitations. First, antibody levels measured at enrolment (week 0) are already elevated above the baseline measured in malaria-naïve control panels. This may be due to a delay in care seeking, or because some individuals in our study have had past exposure to malaria. Second, it is currently impossible to determine whether the symptomatic infections detected at week 0 in our individuals are new infections from mosquito bites or due to relapses from previous infections. The source of infection may play a role in antibody boosting and maintenance. Third, due to the longer interval between sampling (4 weeks), there is the possibility of missed infections present in the asymptomatic cohort, which could contribute to the greater variation observed between individuals, especially towards the end of the study after adjusted for time of infection as the sample size reduced. Additionally, the IgG subclass and IgM profiles were not obtained in the asymptomatic cohort and thus further investigations on IgM responses following *P. vivax* asymptomatic infections will be required. Finally, we only measured total IgG responses in the asymptomatic cohort against the 27 *P. vivax* proteins identified as immunogenic in the symptomatic cohort; the profile against those non-immunogenic proteins may have been different in the individuals with greater past exposure.

In the present study, we have provided new insights that contribute to our understanding of *P. vivax*-specific antibody acquisition and maintenance in low-transmission settings such as western Thailand. Our findings have highlighted the antigen- and time-specific nature of *P. vivax* humoral responses, and these responses could be further assessed in the context of using antibodies as markers of immune status or past exposure [19]. Examination of serological status in populations could facilitate identification of transmission hotspots for timely management and appropriate adjustment of pre-existing intervention programs, or identify populations at risk of resurgence due to waning immunity. From a technical aspect, we have also demonstrated a multiplexed, highly sensitive approach that allows rapid screening of antigenic proteins targeted by antibodies using a minimal amount of plasma. This assay could thus play a role in further advancing the discovery of candidate malaria antigens for various strategies of intervention. Lastly, we show that some newly identified *P. vivax* antigens in our panel can be highly immunogenic and even induce long-lived antibody responses. This finding will broaden our current scope of ‘vaccine candidates’ and facilitate exploration of untraditional targets. Moving forward, investigations on antigenic sequence features and host cellular responses and their associations with antibody maintenance are of high priority. Such information will inform fundamental understandings of immunity in response to malaria infections and therefore contribute to the future development of novel therapeutic or diagnostic tools to further control and accelerate elimination of *P. vivax* malaria and avoid the re-establishment of transmission.

## Materials and Methods

### Study populations

34 symptomatic patients of all ages (see Table 1) with confirmed *P. vivax* infection were enrolled in a 9-month longitudinal study in Tha Song Yang, Tak Province, Thailand, as previously described [52]. These individuals were a sub-set from a larger study [28], and selected due to the absence of recurrent *Plasmodium* infections during follow-up. Treatment was administered according to Thai national guidelines upon confirmation of diagnosis. Plasma samples were collected via finger prick at 17 time points post-infection and treatment: time of infection (w0), w1, w2, then every fortnight for 6 months, then every month until the 9^th^ month. Light microscopy and quantitative *P. vivax*-specific PCR were performed on all samples to confirm the absence of recurrent *Plasmodium* infections during the follow-up.

To compare antibody kinetic profiles with the symptomatic individuals, 30 individuals with asymptomatic *P. vivax* infections were selected from a larger longitudinal cohort study conducted in the Kanchanaburi and Ratchaburi provinces of western Thailand in 2013-2014 [29]. Plasma samples were collected via finger prick at enrolment and approximately every month for a year (a total of 14 visits). The 30 individuals (see Table 1) were detected as *P. vivax*-positive by PCR at early time-points (between enrolment and 6 months) of this yearlong study, and each volunteer had at least 10 samples available post-infection. The infections were classified as asymptomatic as the individuals had no fever (<37.5°C) and no self-reported malaria symptoms at the time of infection.

Plasma samples of 274 malaria-free individuals from the Australian Red Cross (ARC, n=100) in Melbourne, Australia, the Thai Red Cross (TRC, n=72) in Bangkok, Thailand and the Volunteer Blood Donor Registry (VBDR, n=102) in Melbourne, Australia, were collected as previously described [52].

### Antigen selection and expression

Fifty-two *P. vivax* antigens known to be immunogenic in low-transmission regions such as Thailand and Brazil were selected for this study, based on our previous results [24, 52]. Thirty-nine antigens were expressed using a wheat germ cell-free (WGCF) system (CellFree Sciences, Matsuyama, Japan), whilst 13 were expressed using *E. coli*. This panel of antigens included both those that can act as markers of recent exposure [52] and those currently under consideration as vaccine candidates [8, 43, 74-77]. All antigens were expressed and purified as previously described in detail [52]. See Table S2 for the full list of 52 *P. vivax* proteins.

### Antigen conjugation

The conjugation of purified *P. vivax* antigens to non-magnetic COOH microspheres (Bio-Rad Laboratories, Inc., California, USA) was performed as described [52]. The concentration of each antigen used for conjugation was optimised to achieve a standard curve with log-linearity and is recorded in Table S2. In brief, 2.5 × 10^6^ microspheres with unique internal colours were activated before optimal concentration of an antigen was added and incubated either overnight at 4°C or for 2 hours at room temperature in the dark before excess antigen were removed. The standard curve was generated using a positive control plasma pool, as detailed below.

### Total IgG antibody measurements

Total IgG antibody measurements were performed using a multiplexed bead-based assay as previously described [52]. Briefly, 500 antigen-conjugated microspheres per well were added into a MultiScreen Solvinert 96 Well Filter Plate (Merk Millipore) and incubated at room temperature with plasma samples diluted in a 1:100 ratio. Antigen-coupled microspheres were then incubated with a PE-conjugated anti-human secondary antibody at 1:100 dilution (donkey F(ab’)_2_ anti-human IgG Fc, 1 mg/ml, Jackson ImmunoResearch Laboratories, Inc.). For the clinical Tak cohort, IgG levels at all 17 time points were measured. For the asymptomatic Kanchanaburi cohort, IgG levels were measured at time of infection and at a minimum of 10 later time-points. Plasma samples from hyper-immune individuals in Papua New Guinea (PNG) were pooled and used as the positive control from which 2-fold serially-diluted standard curves were established, as described [52].

The assays were performed on either a Bio-Plex 200^®^ (Bio-Rad Laboratories, Inc.) or Luminex^®^ 200™ (Luminex^®^ Corporation, Texas, USA) machine as previously described [78]. MFI values were converted into arbitrary RAU based on the antigen-specific standard curves on each plate, to adjust for any plate-plate variations. The conversion was performed using a 5-parameter logistic regression model [52].

### IgG subclass and IgM antibody measurements

IgM antibody measurements were performed at all 17 time points in the symptomatic cohort and positivity was determined at the peak of response (1-week post-infection). The same multiplexed assay was used as described above for IgG, with the following modifications: samples were diluted at 1:200 and donkey F(ab’)2 anti-human IgM Fc_5μ_ at 1:400 dilution (1 mg/ml, Jackson ImmunoResearch Laboratories, Inc.) was used as the secondary antibody for detection.

To investigate the IgG subclass profile in the symptomatic cohort, measurements were performed at 7 time points: w0, w2, w4, and then every 8 weeks until 9 months. Week 2, rather than peak responses at week 1, was selected due to plasma availability. Again, the multiplexed assay was used, with the following modifications. To maximise signals and minimise non-specific background, samples were diluted at 1:50 and antigen-coupled microspheres were incubated with secondary antibody diluted at 1:100 each: mouse anti-human IgG1 hinge (0.1 mg/ mL, clone 4E3, SouthernBiotech); IgG2 Fc (0.1 mg/ mL, clone HP6002, SouthernBiotech); IgG3 hinge (0.1 mg/ mL clone HP6050, SouthernBiotech); IgG4 Fc (0.1 mg/ mL, clone HP6025, SouthernBiotech).

### Antigen feature selection

To explore the potential factors influencing estimated antibody half-lives, 6 antigen features were selected for analysis: signal peptides, transmembrane domains, GPI anchors, PEXEL motifs, predicted domains and low-complexity regions. The presence of these features in a sequence was as deposited in PlasmoDB [79] (https://plasmodb.org/plasmo/app) or as predicted using the online servers: Transmembrane Helices Hidden Markov Model (TMHMM) Server version 2.0 [80] (http://www.cbs.dtu.dk/services/TMHMM/), PredGPI [81] (http://gpcr.biocomp.unibo.it/predgpi/pred.htm), InterPro [82] (https://www.ebi.ac.uk/interpro/), and SEG [83].

### Mathematical modelling

Following infection, we assume that the proliferation and differentiation of B cells leads to a boost in antibody secreting cells (ASC) of size β, commencing at a time *δ* before the first sample. A proportion *ρ* of these ASCs are assumed to be short-lived with half-life *d*_*s*_, with a proportion 1 – *ρ* being long-lived with half-life *d*_*l*_. It is assumed that all ASCs secrete IgG molecules which decay with a half-life *d*_*a*_. The antibody level of an individual at time *t* after first sample is given by:

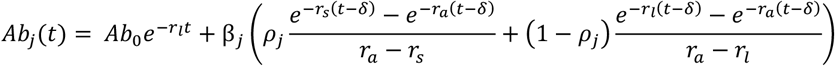

where *r*_*a*_ = log(2)/*d*_*a*_ is the rate of decay of IgG molecules, *r*_*s*_ = log(2)/*d*_*s*_ is the rate of decay of short-lived plasma B cells, and *r*_*l*_ = log(2)/*d*_*l*_ is the rate of decay of long-lived plasma B cells. It is assumed that the measured antibody level before vaccination was *Ab*_*0*_ which decays exponentially at rate *r*_*l*_.

The model was fitted to longitudinal antibody level measurements from all participants. Mixed effects methods were used to capture the natural variation in antibody kinetics between individual participants, whilst estimating the average value and variance of the immune parameters across the entire population of individuals. The models were fitted in a Bayesian framework using Markov Chain Monte Carlo (MCMC) methods. Mixed effects methods allow individual-level parameters to be estimated for each participant separately, with these individual-level (or mixed effects) parameters being drawn from global distributions. For example, for each participant *n* the half-life of the short-lived ASCs may be estimated as 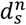 (an individual-level parameter). These *N* estimates of the local parameters 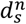 will be drawn from a probability distribution. A log-Normal distribution is suitable as it has positive support on [0, ∞). Thus we have 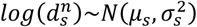. The mean *d*_*s*_ and the variance 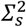 of the estimates of 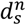 are given by 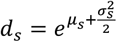 and 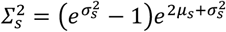. The relationship between the parameters describing the population-level distribution and the parameters for each individual in the population are depicted in the schematic diagram in Figure S8.

The model likelihood was calculated via the following process. For individual *n* we have data on observed antibody levels *A*^*n*^ = {*a*_1_, …, *a*_*K*_} at times *T*^*n*^ = {*t*_1_, …, *t*_*K*_}. We denote *D*^*n*^ = (*A*^*n*^, *T*^*n*^) to be the vector of data for individual *n*. For individual *n*, the parameters 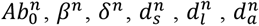 and *ρ*^*n*^ are estimated. These parameters are denoted 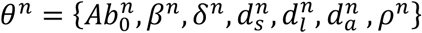. The model predicted antibody levels will be {*A*(*t*_1_), *A*(*t*_2_), …, *A*(*t*_*K*_)}. We assume log-Normally distributed measurement error such that the difference between log(*a*_*j*_) and log(*Ab*(*t*_*j*_)) is Normally distributed with variance 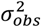. For model predicted antibody levels *Ab*(*t*_*j*_) the data likelihood for individual *n* is given by

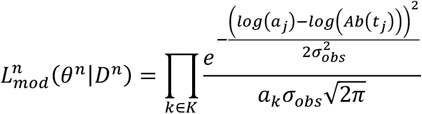

As described above, for each individual there are 7 parameters to be estimated: 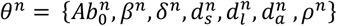. The mixed effects likelihood can be written as follows:

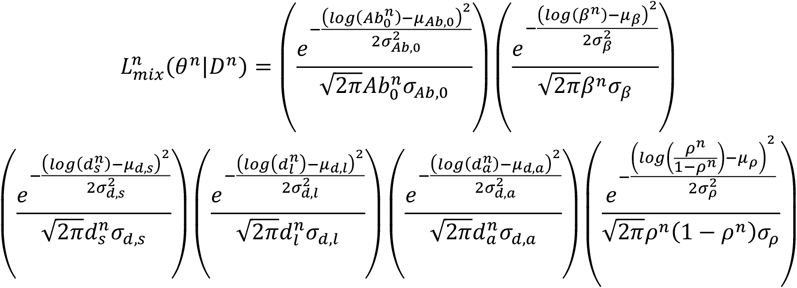

As the proportion of the ASCs that are long-lived must be bounded by 0 and 1, the individual-level parameters *ρ*^*n*^ are assumed to be drawn from logit-Normal distributions. Note that the pre-existing antibody level 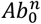 will be variable, depending on a large number of covariates such as age and past exposure. We therefore do not attempt to constrain pre-existing antibody levels using mixed effects.

Denote *D* = {*D*^1^, …, *D*^*N*^} to be the vector of data for all *N* participants. We denote θ to be the combined vector of population-level parameters and individual-level parameters to be estimated. The total likelihood is obtained by multiplying the likelihood for each participant

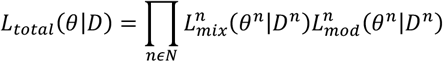

The model was fitted to the data using Markov Chain Monte Carlo (MCMC) methods using software for Bayesian statistical inference of non-linear mixed-effects models. This utilises a Metropolis-within-Gibbs sampler, whereby population-level parameters are updated using a Gibbs sampler, and individual-level and observational parameters are updated using a Metropolis-Hastings sampler.

### Statistical analyses

Statistical tests, including Fisher’s exact test, Mann-Whitney test and Spearman’s correlation test, were performed using R version 4.0.4 and GraphPad Prism version 7.0. Linear and logistic regression models to establish associations between antigen features and antibody responses were performed using Stata version 12.1.

### Ethics statement

Collection of the Thai symptomatic longitudinal study in Tha Song Yang, Tak Province, was approved by The Ethics Committee of the Faculty of Tropical Medicine, Mahidol University, Thailand (MUTM 2014-025-01 and -02). For the Thai asymptomatic cohort study in the Kanchanaburi and Ratchaburi provinces, collection of samples was approved by The Ethics Committee of the Faculty of Tropical Medicine, Mahidol University, Thailand (MUTM 2013-027-01). The Human Research Ethics Committee (HREC) at the Walter and Eliza Hall Institute of Medical Research (WEHI) approved the use of these samples in Melbourne (#14/02). The collection and/or use of the 3 malaria-free control panels in Melbourne were approved by the HREC at WEHI (#14/02).

## Acknowledgments

We gratefully acknowledge the extensive field teams that contributed to sample collection and qPCR assays in the original cohort studies. We thank all of the individuals who participated in each of the studies, and the Australian and Thai Red Cross for donation of blood samples. We thank the Volunteer Blood Donor Registry at WEHI for donation of plasma samples and Lina Laskos for their collection and advice. We thank Christopher King (Case Western Reserve University) for provision of the PNG control plasma pool. We thank Fumie Matsuura (CellFree Sciences), Christele Huon (Institut Pasteur) and Jakub Gruszczyk (WEHI) for contributing to expression of proteins. We thank Connie Li-Wai-Suen for writing the R script used for converting MFI values to relative antibody units.

## Supplementary Figures and Tables

**Figure S1.**
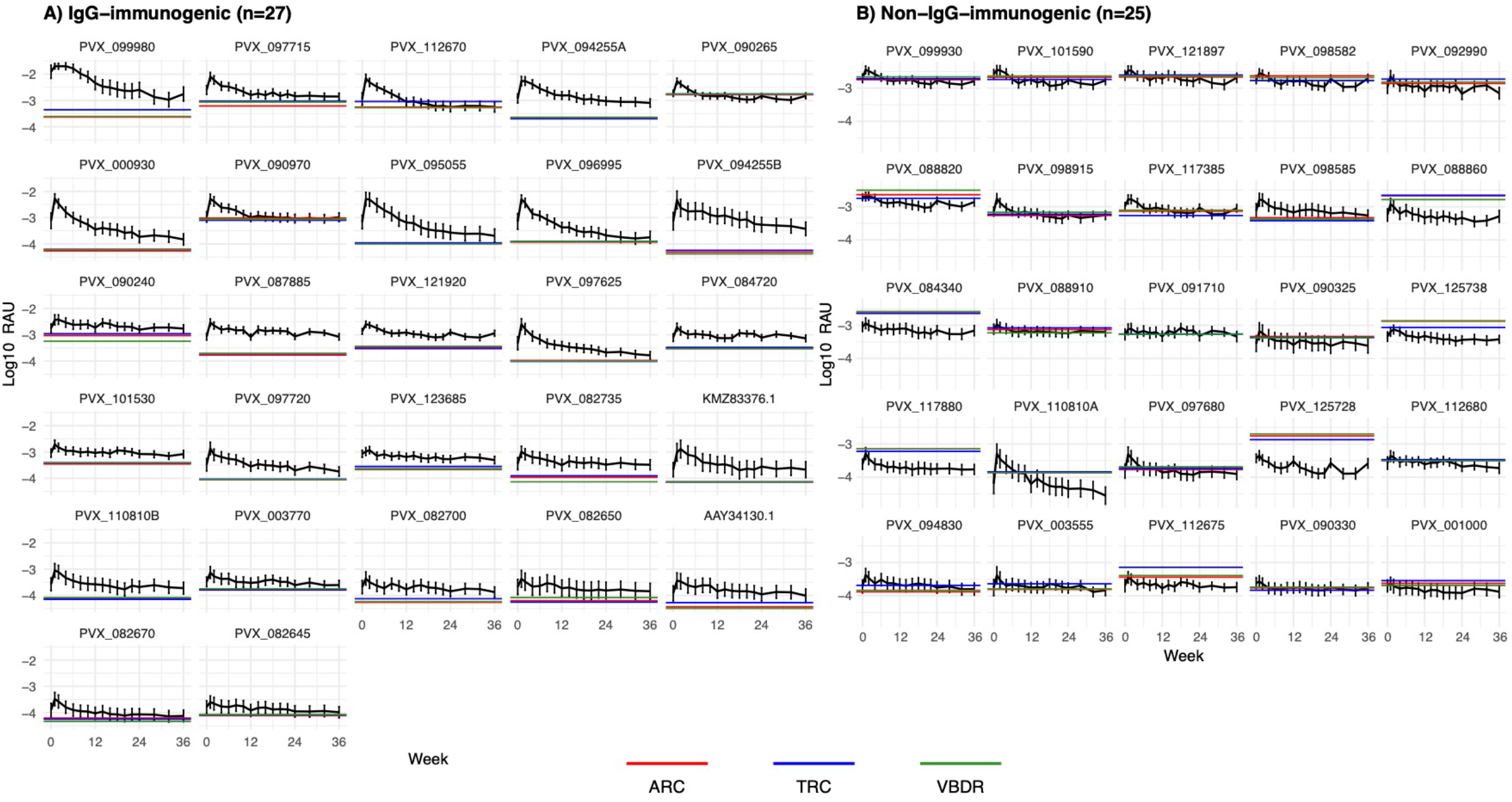
Total IgG kinetic profiles against 52 *P. vivax* antigens in 34 Thai patients with symptomatic *P. vivax* infections. Antigen-specific IgG antibody responses were measured for 9 months following a symptomatic *P. vivax* infection among 34 patients from western Thailand. The observed kinetic profiles were categorised into **A)** IgG-immunogenic (> 1 SD at 1-week post-infection above negative control baseline; n = 27) and **B)** non-IgG-immunogenic (> 1 SD at 1-week post-infection; n = 25). The horizontal lines represent the negative control panels which consist of malaria-free individuals from 3 sources: Australian Red Cross (ARC; n = 100), Thai Red Cross (TRC; n = 72) and Volunteer Blood Donor Registry (VBDR; n = 102). Data are expressed as median ± 95% CI of the 34 patients.

**Figure S2.**
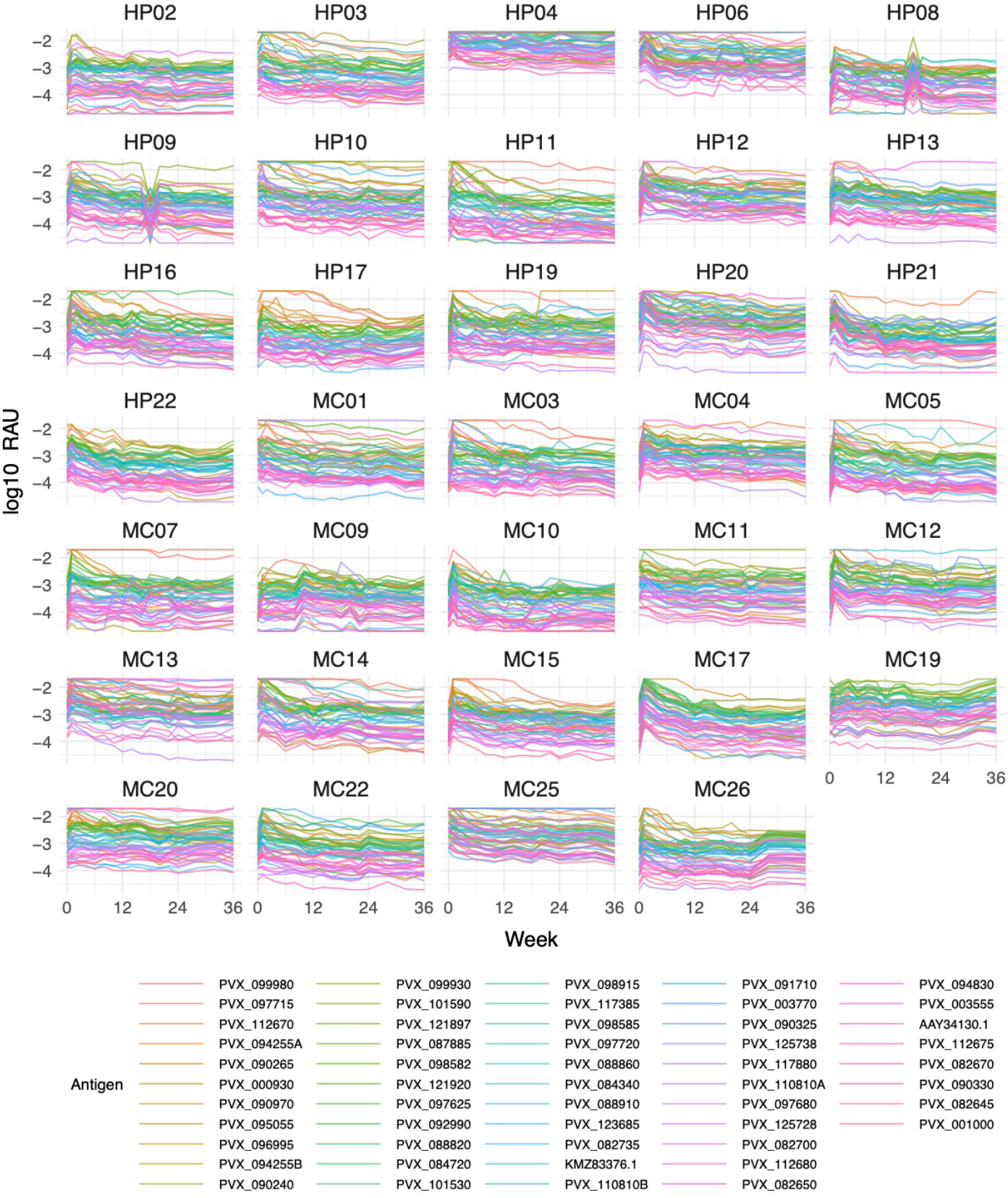
Adjusted total IgG kinetic profiles at an individual level following symptomatic infections. Antigen-specific IgG antibody responses were measured for 9 months following a symptomatic *P. vivax* infection among 34 patients from western Thailand. The responses of each of the 34 participants were plotted per *P. vivax* antigen. Note that a limitation in this data is the peak for HP08 and drop for HP09 at week 18; we assume that a labelling error occurred in the field and that these two samples are swapped, however we have not altered the data to account for this.

**Table S1.** Immunogenicity of total IgG against 52 *P. vivax* antigens at 1-week post-infection in Thai symptomatic volunteers.

| Protein ID<sup>a</sup> | Median<sup>b</sup> | Number of volunteers (%) |  |  |  |  |  |  |
| --- | --- | --- | --- | --- | --- | --- | --- | --- |
|  |  | 95% CI<sup>b</sup> |  | 1% of | 5% of | 10% of | 25% of | 50% of |
|  |  |  |  | PNG levels | PNG levels | PNG levels | PNG levels | PNG levels |
| PVX_099980 | 20.00 | 9.15 | 20.00 | 34 (1.0) | 34 (1.0) | 34 (1.0) | 34 (1.0) | 17 (0.5) |
| PVX_096995 | 4.99 | 2.40 | 15.29 | 34 (1.0) | 34 (1.0) | 33 (1.0) | 27 (0.8) | 20 (0.6) |
| PVX_088860 | 1.18 | 0.63 | 15.29 | 34 (1.0) | 28 (0.8) | 19 (0.6) | 8 (0.2) | 7 (0.2) |
| PVX_101530 | 1.95 | 0.84 | 3.28 | 34 (1.0) | 32 (0.9) | 23 (0.7) | 15 (0.4) | 8 (0.2) |
| PVX_112680 | 0.43 | 0.26 | 0.97 | 34 (1.0) | 20 (0.6) | 11 (0.3) | 5 (0.1) | 3 (0.1) |
| PVX_097715 | 8.13 | 3.37 | 19.14 | 34 (1.0) | 34 (1.0) | 32 (0.9) | 29 (0.9) | 22 (0.6) |
| PVX_094830 | 0.42 | 0.20 | 1.41 | 34 (1.0) | 18 (0.5) | 13 (0.4) | 8 (0.2) | 2 (0.1) |
| PVX_112675 | 0.36 | 0.25 | 0.52 | 32 (0.9) | 15 (0.4) | 8 (0.2) | 5 (0.1) | 2 (0.1) |
| PVX_112670 | 6.97 | 2.50 | 15.91 | 34 (1.0) | 34 (1.0) | 32 (0.9) | 28 (0.8) | 23 (0.7) |
| PVX_090970 | 5.14 | 1.61 | 13.60 | 34 (1.0) | 32 (0.9) | 29 (0.9) | 24 (0.7) | 18 (0.5) |
| PVX_084720 | 1.94 | 1.04 | 4.07 | 34 (1.0) | 33 (1.0) | 25 (0.7) | 12 (0.4) | 7 (0.2) |
| PVX_003770 | 0.75 | 0.27 | 2.94 | 34 (1.0) | 24 (0.7) | 18 (0.5) | 12 (0.4) | 8 (0.2) |
| PVX_092990 | 2.27 | 1.06 | 5.70 | 34 (1.0) | 33 (1.0) | 29 (0.9) | 22 (0.6) | 11 (0.3) |
| PVX_091710 | 0.76 | 0.59 | 2.29 | 34 (1.0) | 26 (0.8) | 17 (0.5) | 9 (0.3) | 7 (0.2) |
| PVX_087885 | 2.93 | 1.51 | 5.22 | 34 (1.0) | 33 (1.0) | 29 (0.9) | 19 (0.6) | 9 (0.3) |
| PVX_003555 | 0.41 | 0.16 | 1.37 | 34 (1.0) | 20 (0.6) | 15 (0.4) | 9 (0.3) | 6 (0.2) |
| PVX_117385 | 1.65 | 1.08 | 3.88 | 34 (1.0) | 31 (0.9) | 22 (0.6) | 11 (0.3) | 5 (0.1) |
| PVX_090265 | 5.37 | 2.59 | 7.43 | 34 (1.0) | 34 (1.0) | 31 (0.9) | 25 (0.7) | 17 (0.5) |
| PVX_082700 | 0.44 | 0.19 | 2.39 | 27 (0.8) | 15 (0.4) | 12 (0.4) | 10 (0.3) | 7 (0.2) |
| PVX_082650 | 0.43 | 0.12 | 5.06 | 27 (0.8) | 17 (0.5) | 12 (0.4) | 9 (0.3) | 9 (0.3) |
| PVX_094255A<sup>c</sup> | 5.46 | 2.45 | 7.84 | 34 (1.0) | 34 (1.0) | 29 (0.9) | 29 (0.9) | 21 (0.6) |
| PVX_097680 | 0.49 | 0.16 | 1.39 | 33 (1.0) | 19 (0.6) | 10 (0.3) | 7 (0.2) | 5 (0.1) |
| PVX_001000 | 0.23 | 0.14 | 0.80 | 29 (0.9) | 14 (0.4) | 8 (0.2) | 5 (0.1) | 2 (0.1) |
| PVX_097625 | 2.52 | 0.85 | 5.16 | 34 (1.0) | 30 (0.9) | 28 (0.8) | 19 (0.6) | 12 (0.4) |
| PVX_082670 | 0.33 | 0.14 | 1.69 | 25 (0.7) | 12 (0.4) | 10 (0.3) | 8 (0.2) | 6 (0.2) |
| PVX_099930 | 3.53 | 2.40 | 6.53 | 34 (1.0) | 34 (1.0) | 33 (1.0) | 19 (0.6) | 9 (0.3) |
| PVX_084340 | 1.15 | 0.78 | 4.09 | 34 (1.0) | 27 (0.8) | 17 (0.5) | 10 (0.3) | 7 (0.2) |
| PVX_098915 | 1.75 | 0.89 | 2.24 | 34 (1.0) | 29 (0.9) | 23 (0.7) | 8 (0.2) | 6 (0.2) |
| PVX_088820 | 2.01 | 1.55 | 3.47 | 34 (1.0) | 34 (1.0) | 31 (0.9) | 13 (0.4) | 7 (0.2) |
| PVX_082735 | 1.02 | 0.41 | 2.20 | 32 (0.9) | 21 (0.6) | 16 (0.5) | 10 (0.3) | 7 (0.2) |
| PVX_082645 | 0.25 | 0.15 | 1.69 | 28 (0.8) | 13 (0.4) | 10 (0.3) | 7 (0.2) | 3 (0.1) |
| PVX_117880 | 0.52 | 0.27 | 1.85 | 33 (1.0) | 17 (0.5) | 14 (0.4) | 7 (0.2) | 3 (0.1) |
| PVX_121897 | 3.37 | 2.03 | 7.57 | 34 (1.0) | 34 (1.0) | 30 (0.9) | 20 (0.6) | 13 (0.4) |
| PVX_125728 | 0.45 | 0.25 | 0.71 | 34 (1.0) | 21 (0.6) | 10 (0.3) | 2 (0.1) | 1 (0.0) |
| PVX_090330 | 0.27 | 0.17 | 0.92 | 27 (0.8) | 12 (0.4) | 9 (0.3) | 4 (0.1) | 1 (0.0) |
| PVX_123685 | 1.03 | 0.54 | 2.29 | 34 (1.0) | 24 (0.7) | 13 (0.4) | 9 (0.3) | 3 (0.1) |
| PVX_125738 | 0.65 | 0.41 | 1.83 | 34 (1.0) | 13 (0.4) | 10 (0.3) | 4 (0.1) | 2 (0.1) |
| PVX_097720 | 1.37 | 0.31 | 3.50 | 31 (0.9) | 22 (0.6) | 19 (0.6) | 13 (0.4) | 7 (0.2) |
| PVX_000930 | 5.29 | 1.66 | 11.96 | 34 (1.0) | 34 (1.0) | 32 (0.9) | 25 (0.7) | 20 (0.6) |

| Protein ID<sup>a</sup> | Median<sup>b</sup> | 95% CI<sup>b</sup> |  | Number of volunteers (%) |  |  |  |  |
| --- | --- | --- | --- | --- | --- | --- | --- | --- |
|  |  |  |  | 1% of<br>PNG levels | 5% of<br>PNG levels | 10% of<br>PNG levels | 25% of<br>PNG levels | 50% of<br>PNG levels |
| PVX_098585 | 1.62 | 0.78 | 4.39 | 34 (1.0) | 30 (0.9) | 25 (0.7) | 15 (0.4) | 10 (0.3) |
| PVX_098582 | 2.81 | 1.91 | 4.56 | 34 (1.0) | 34 (1.0) | 31 (0.9) | 14 (0.4) | 7 (0.2) |
| PVX_121920 | 2.54 | 1.69 | 3.55 | 34 (1.0) | 34 (1.0) | 30 (0.9) | 15 (0.4) | 4 (0.1) |
| PVX_094255B<sup>c</sup> | 4.72 | 1.20 | 14.10 | 29 (0.9) | 28 (0.8) | 26 (0.8) | 25 (0.7) | 22 (0.6) |
| PVX_090325 | 0.67 | 0.21 | 2.82 | 34 (1.0) | 19 (0.6) | 15 (0.4) | 9 (0.3) | 6 (0.2) |
| PVX_101590 | 3.47 | 2.09 | 8.45 | 34 (1.0) | 34 (1.0) | 30 (0.9) | 20 (0.6) | 12 (0.4) |
| PVX_110810A<sup>c</sup> | 0.51 | 0.11 | 1.46 | 29 (0.9) | 21 (0.6) | 15 (0.4) | 9 (0.3) | 6 (0.2) |
| PVX_090240 | 3.70 | 1.10 | 15.50 | 34 (1.0) | 34 (1.0) | 29 (0.9) | 19 (0.6) | 15 (0.4) |
| PVX_088910 | 1.07 | 0.67 | 1.62 | 34 (1.0) | 21 (0.6) | 15 (0.4) | 5 (0.1) | 3 (0.1) |
| PVX_095055 | 5.10 | 1.18 | 17.45 | 34 (1.0) | 30 (0.9) | 28 (0.8) | 21 (0.6) | 18 (0.5) |
| PVX_110810B<sup>c</sup> | 0.90 | 0.35 | 3.18 | 33 (1.0) | 25 (0.7) | 19 (0.6) | 11 (0.3) | 8 (0.2) |
| AAY34130.1<sup>b</sup> | 0.37 | 0.15 | 2.49 | 30 (0.9) | 19 (0.6) | 12 (0.4) | 10 (0.3) | 5 (0.1) |
| KMZ83376.1<sup>b</sup> | 0.97 | 0.11 | 5.04 | 28 (0.8) | 21 (0.6) | 18 (0.5) | 16 (0.5) | 11 (0.3) |

<sup>a</sup> PlasmoDB codes (http://plasmodb.org/plasmo/), <sup>b</sup> Total IgG levels are expressed in relative antibody units (RAU) interpolated from standard curves using a 5PL logistic regression model. Values were multiplied by 1000, <sup>c</sup> A and B represent two distinct protein fragments derived from one full-length protein sequence and hence are expressed by identical PlasmoDB code. Abbreviations: 95% CI = 95% confidence interval

**Figure S3.**
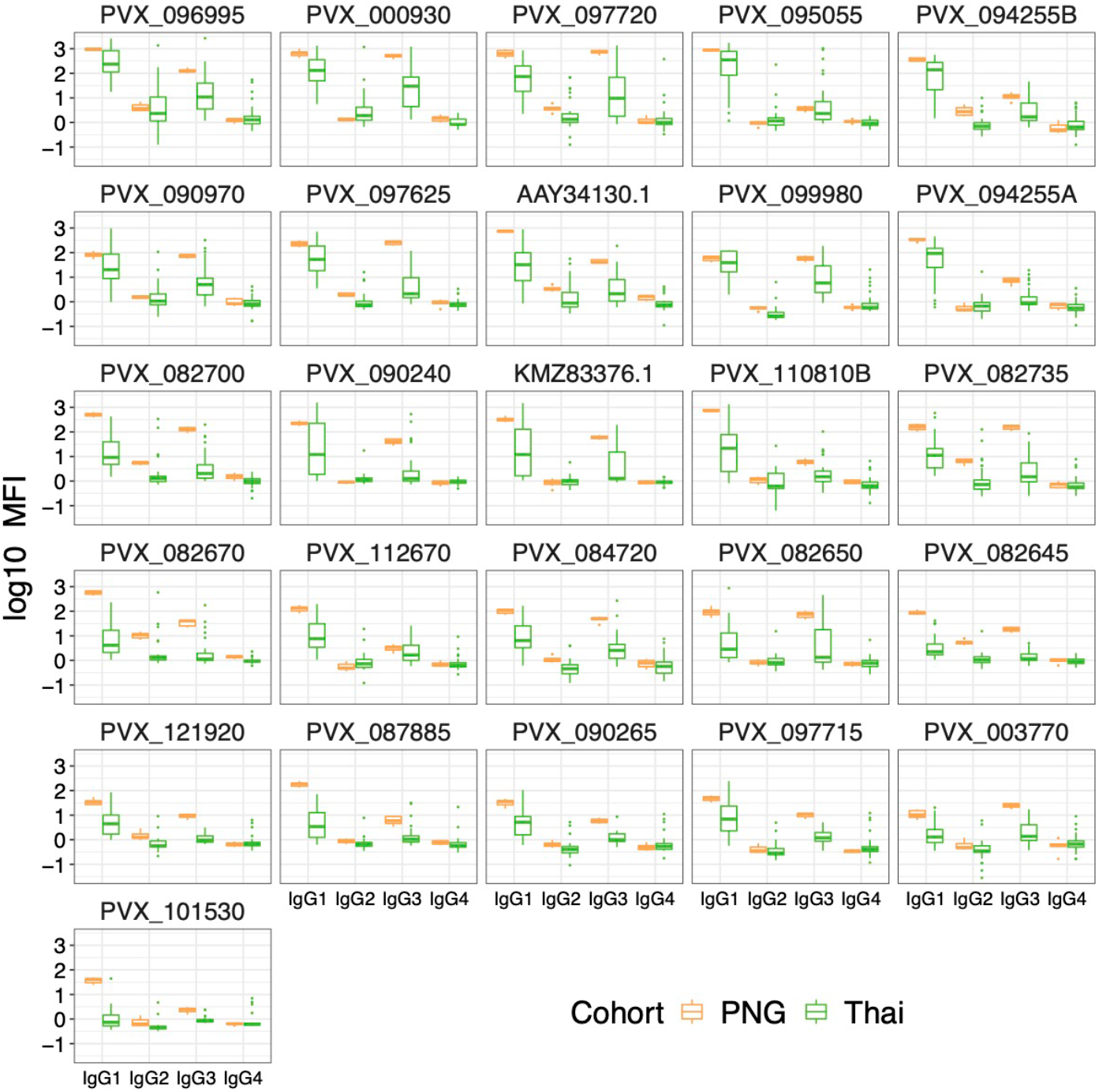
Comparison of the adjusted peak IgG subclass magnitude between symptomatic Thai patients and the PNG hyperimmune pool. Antigen-specific IgG subclass responses against 26 IgG-immunogenic (PVX_123685 was excluded) antigens were measured for 9 months, and the peak of responses (2-week post-infection) was compared between symptomatic Thai patients and the positive control (hyper-immune pooled plasma of infected PNG individuals). PNG results were collected from the standard curves included in each independent experiment of IgG subclass measurement (n=7). Data were adjusted by subtracting the median of pooled negative control panels (n=274).

**Figure S4.**
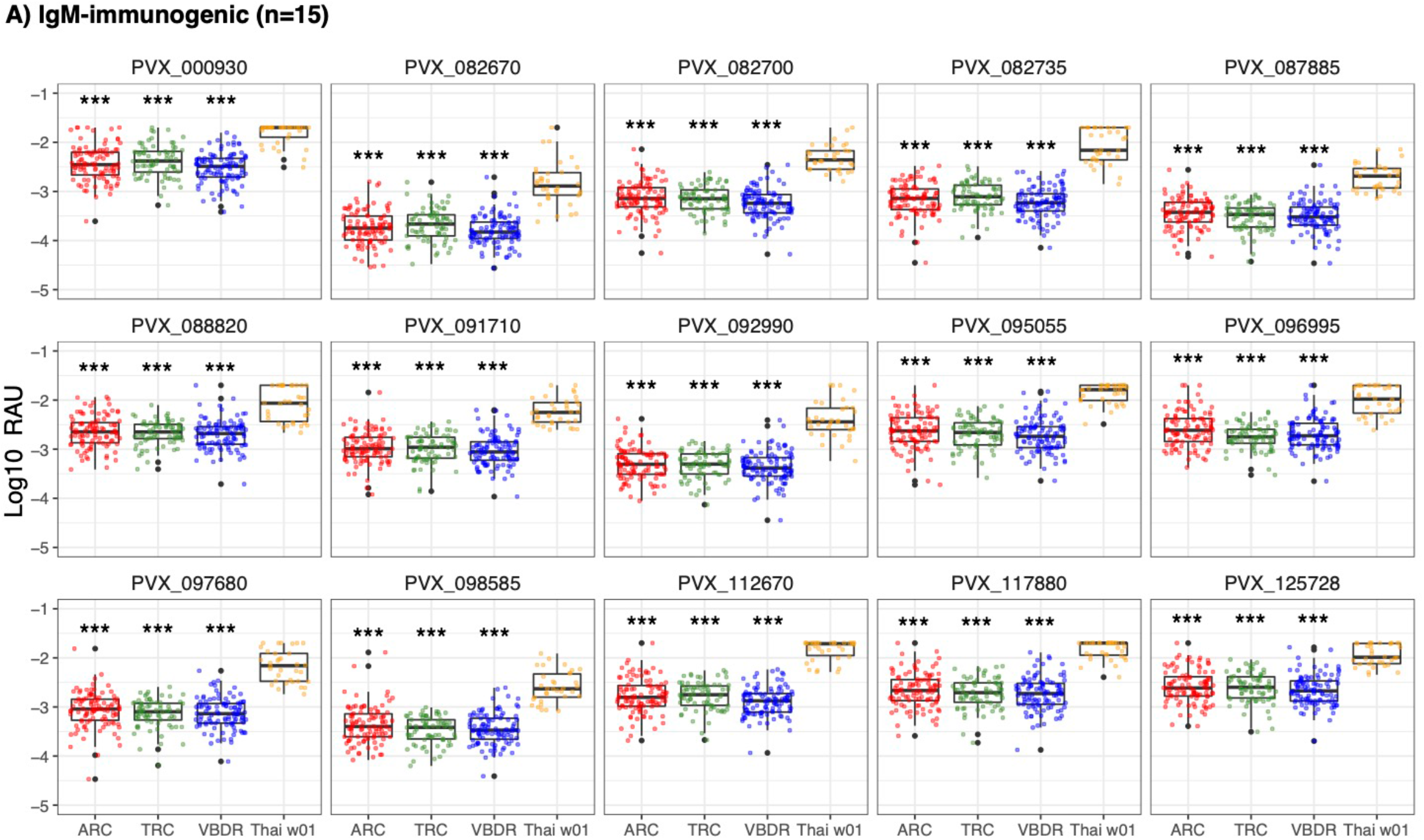
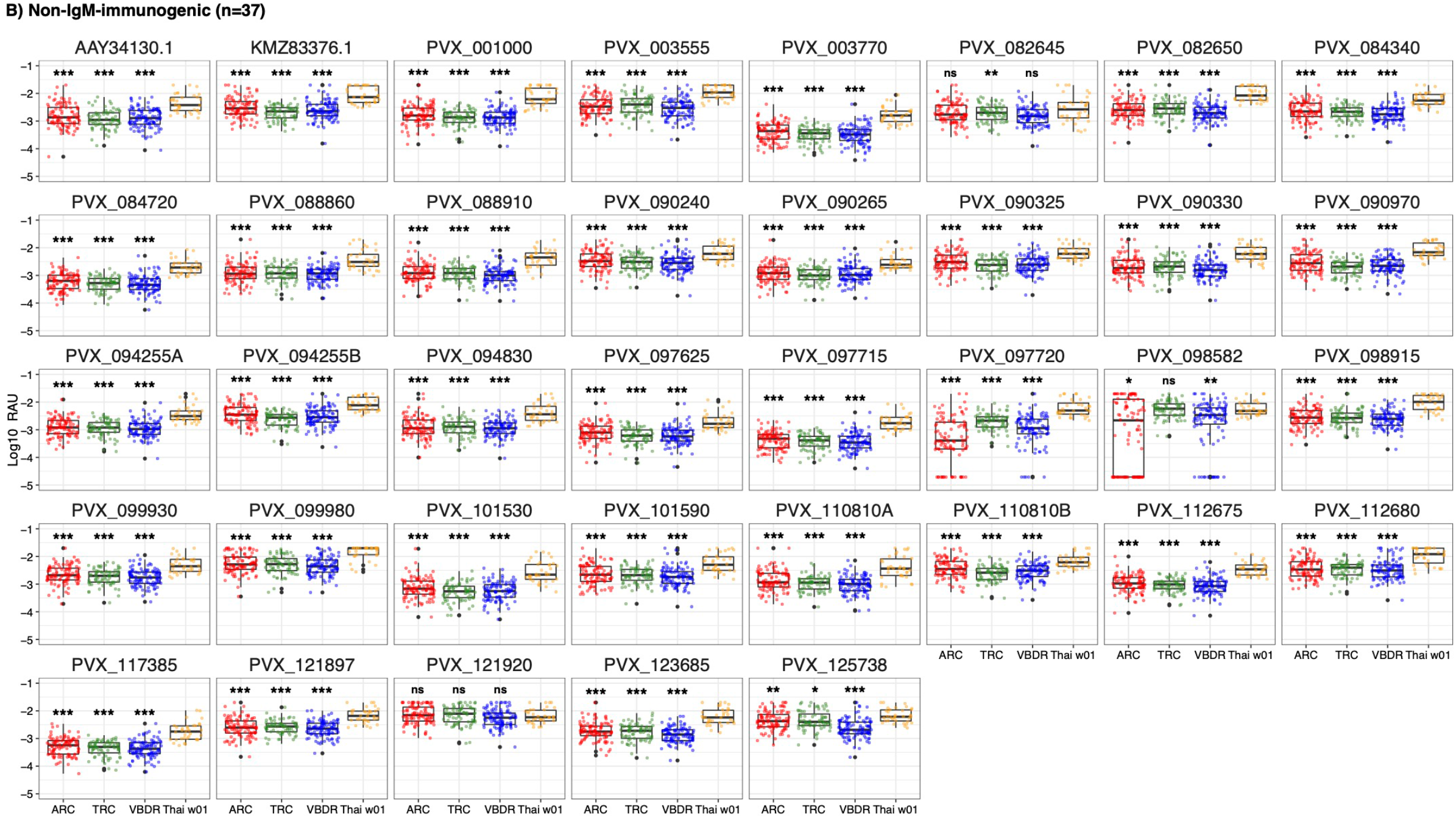
Comparison of the peak IgM magnitude between symptomatic Thai patients and malaria-free individuals. IgM level was measured at the peak of responses (1 week following a symptomatic *P. vivax* infection) among 34 Thai patients and was compared to ARC (n = 100), TRC (n = 72) and VBDR (n = 102). Mann-Whitney’s (non-parametric) test was performed (p > 0.05 ns, < 0.05 *, < 0.01 **, < 0.001 ***). To down-select IgM-positive antigens, medians and standard deviations (SD) were calculated and plotted to identify **A)** antigens with medians more than 2 SD above the pooled negative control panels (n=15) and **B)** those without (n=37).

**Figure S5.**
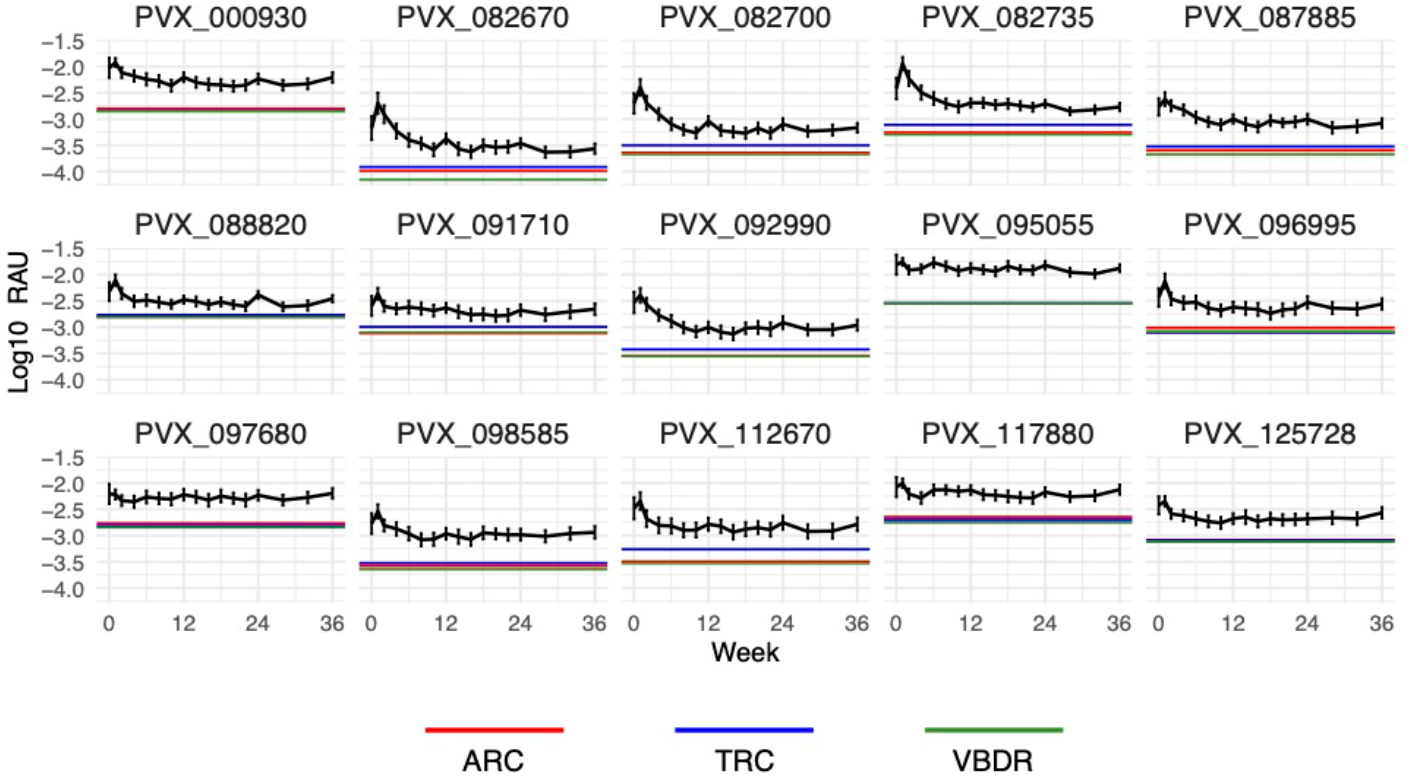
Unadjusted IgM kinetics against 15 *P. vivax* antigens in 34 Thai individuals following symptomatic *P. vivax* infections. 15 *P. vivax* antigens with IgM positivity (> 2 SD above baseline median) at 1-week post-infection were selected and IgM responses over a 9-month period were measured in symptomatic Thai individuals (n=17 timepoints). Horizontal solid lines represent the baseline responses measured in malaria-naïve individuals from 3 different sources (n=274). Data are expressed as the median ± 95% CI.

**Table S2.**
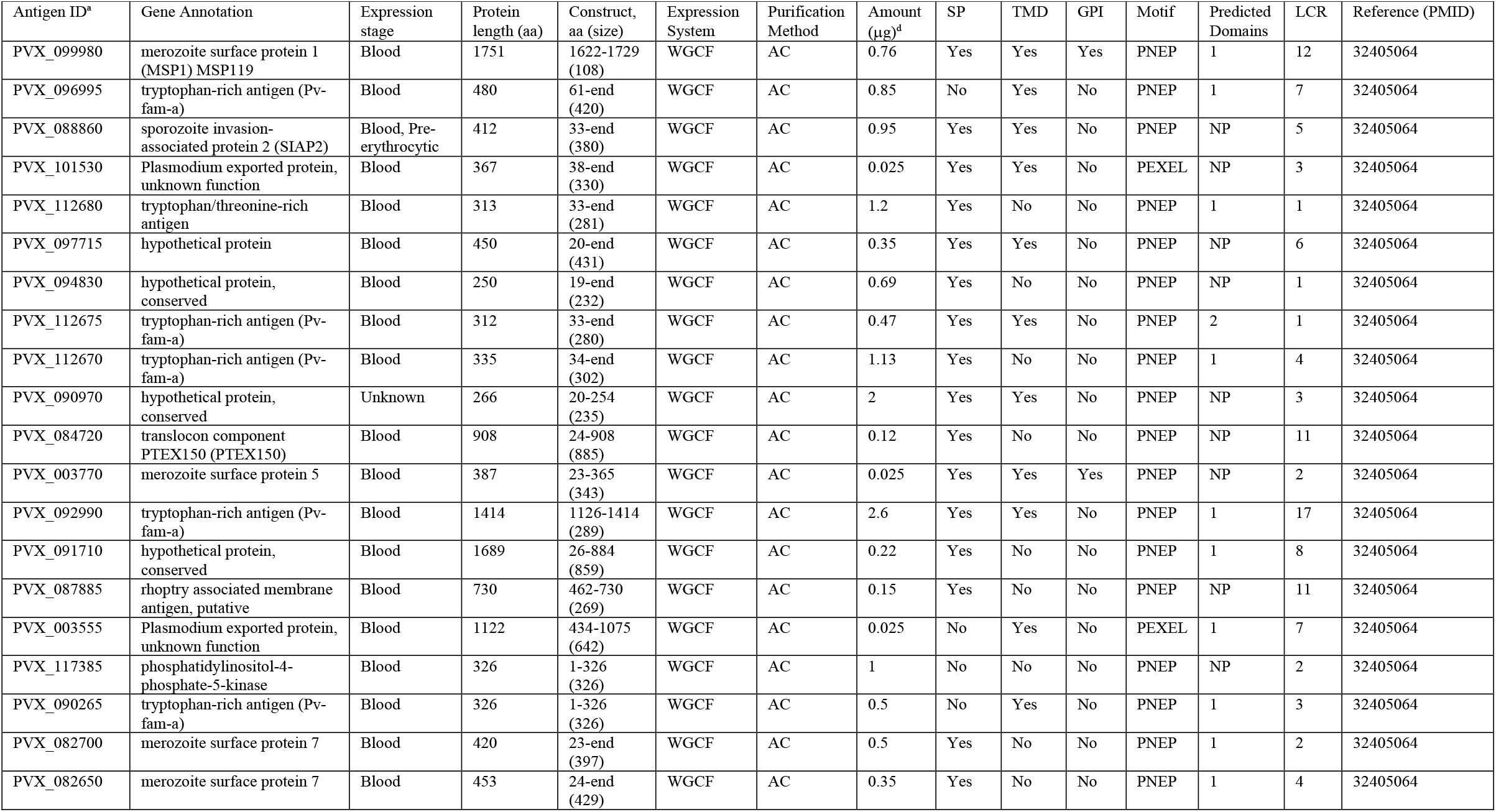
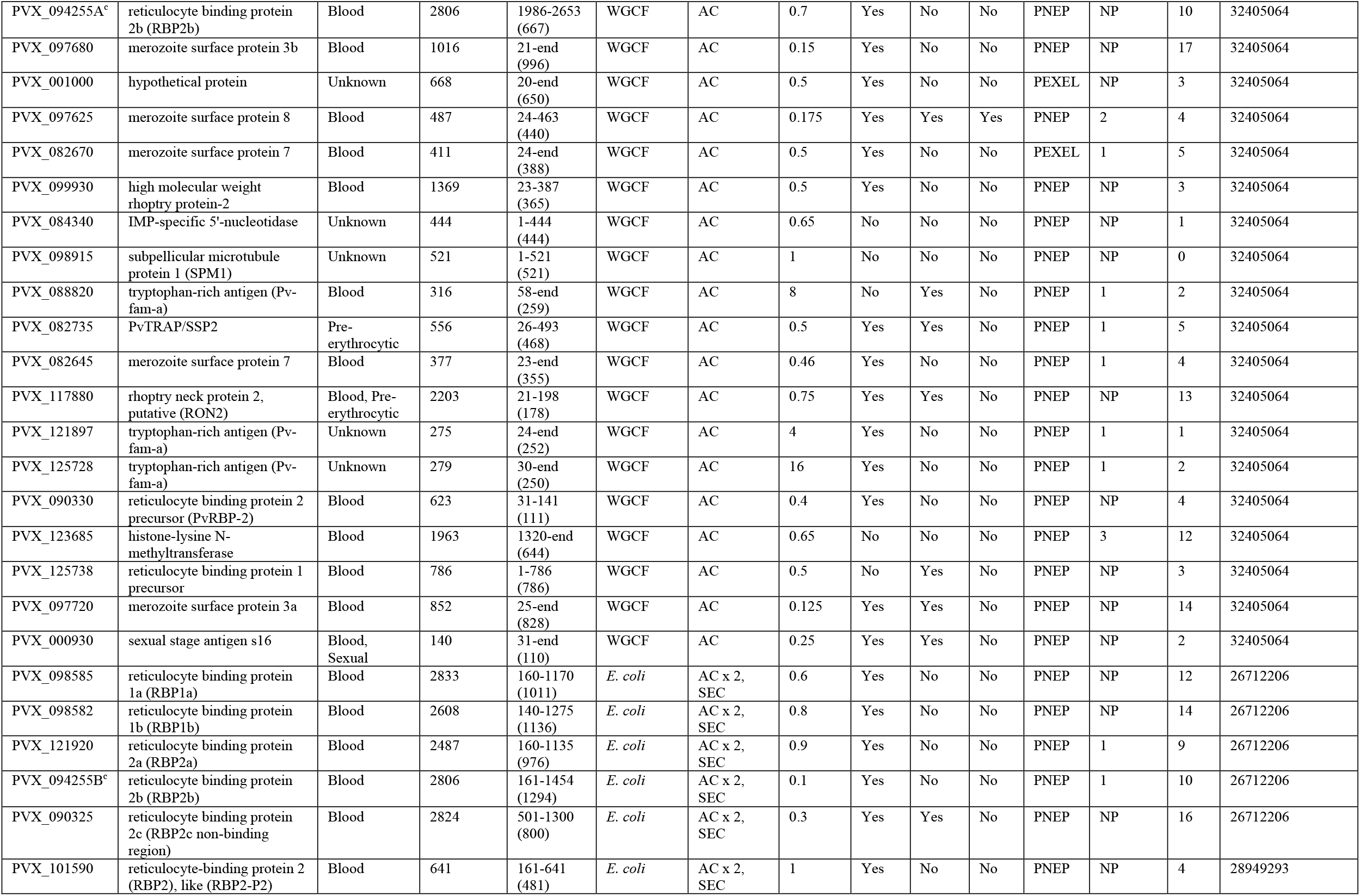
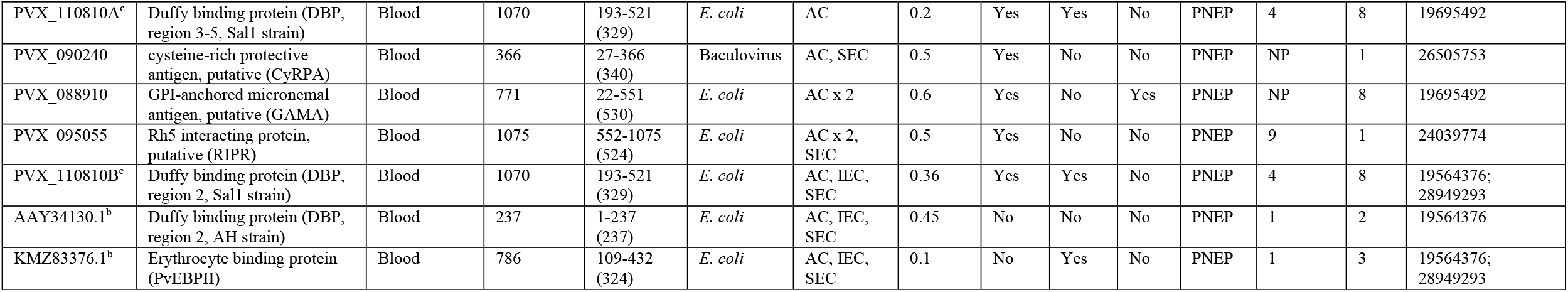
*P. vivax* proteins used in this study. ^a^PlasmoDB codes (http://plasmodb.org/plasmo/), ^b^GenBank IDs, ^c^A and B represent two distinct protein fragments derived from one full-length protein sequence and hence are expressed by identical PlasmoDB code, ^d^Amount refers to the amount of antigen coupled to non-magnetic microspheres. Abbreviations: aa = Amino acid, WGCF = Wheat germ cell-free, E. coli = Escherichia coli, AC = Affinity chromatography, SEC = Size exclusion chromatography, IEC = Ion exchange chromatography, SP = Signal peptide, TMD = transmembrane domain, GPI = Glycosylphosphatidylinositol anchors, PEXEL = Plasmodium exported element, PNEP = PEXEL-negative exported protein, NP = Non-predicted, LCR = Low complexity region.

**Figure S6.**
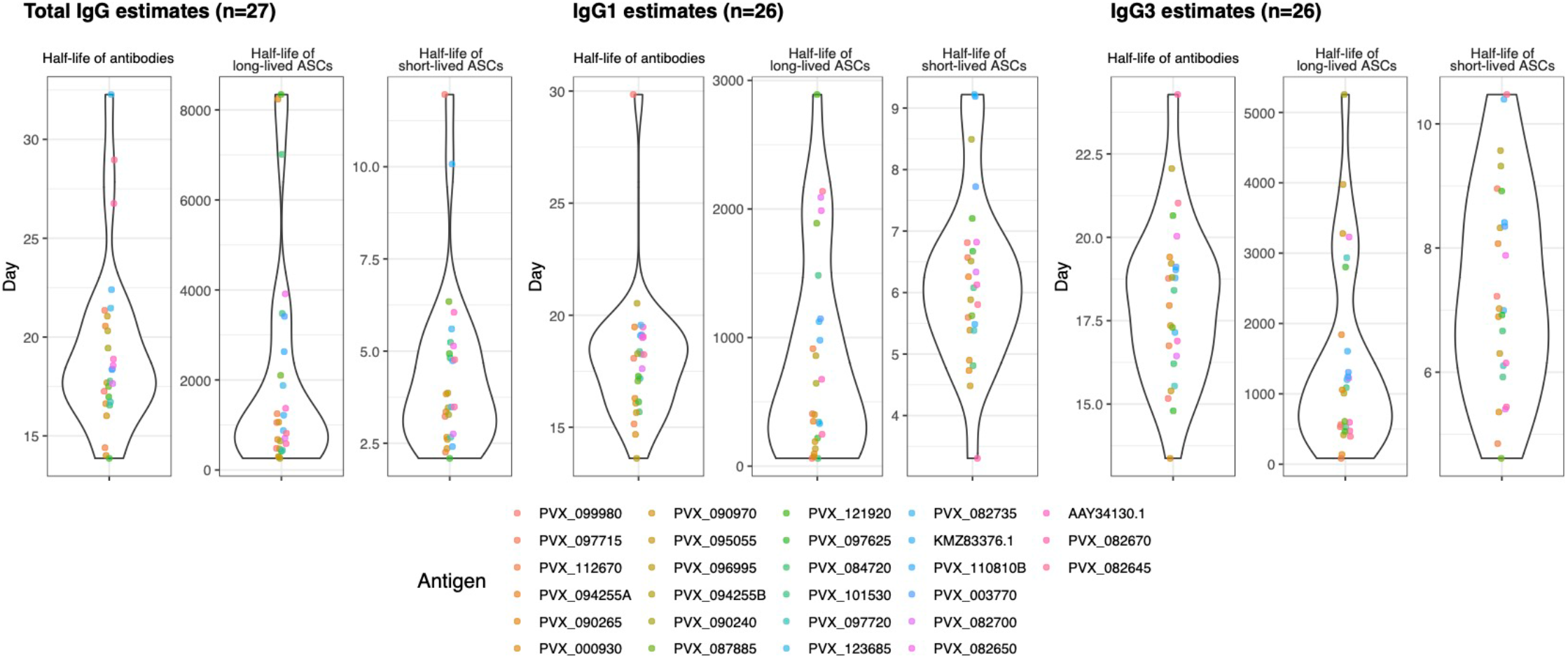
Distribution of data for estimated half-lives of antibodies and ASCs against IgG-immunogenic antigens following symptomatic *P. vivax* infections in the Thai symptomatic cohort. The kinetics of total IgG and IgG subclass antibodies against 52 *P. vivax* antigens were determined following symptomatic *P. vivax* infections in Thai individuals in the absence of recurrent infections for 9 months. Mathematical modelling was then used to generate the estimated half-life of antibodies and ASCs for total IgG, IgG1 and IgG3 per antigen. The difference in distribution between antibody isotypes was visualised.

**Figure S7.**
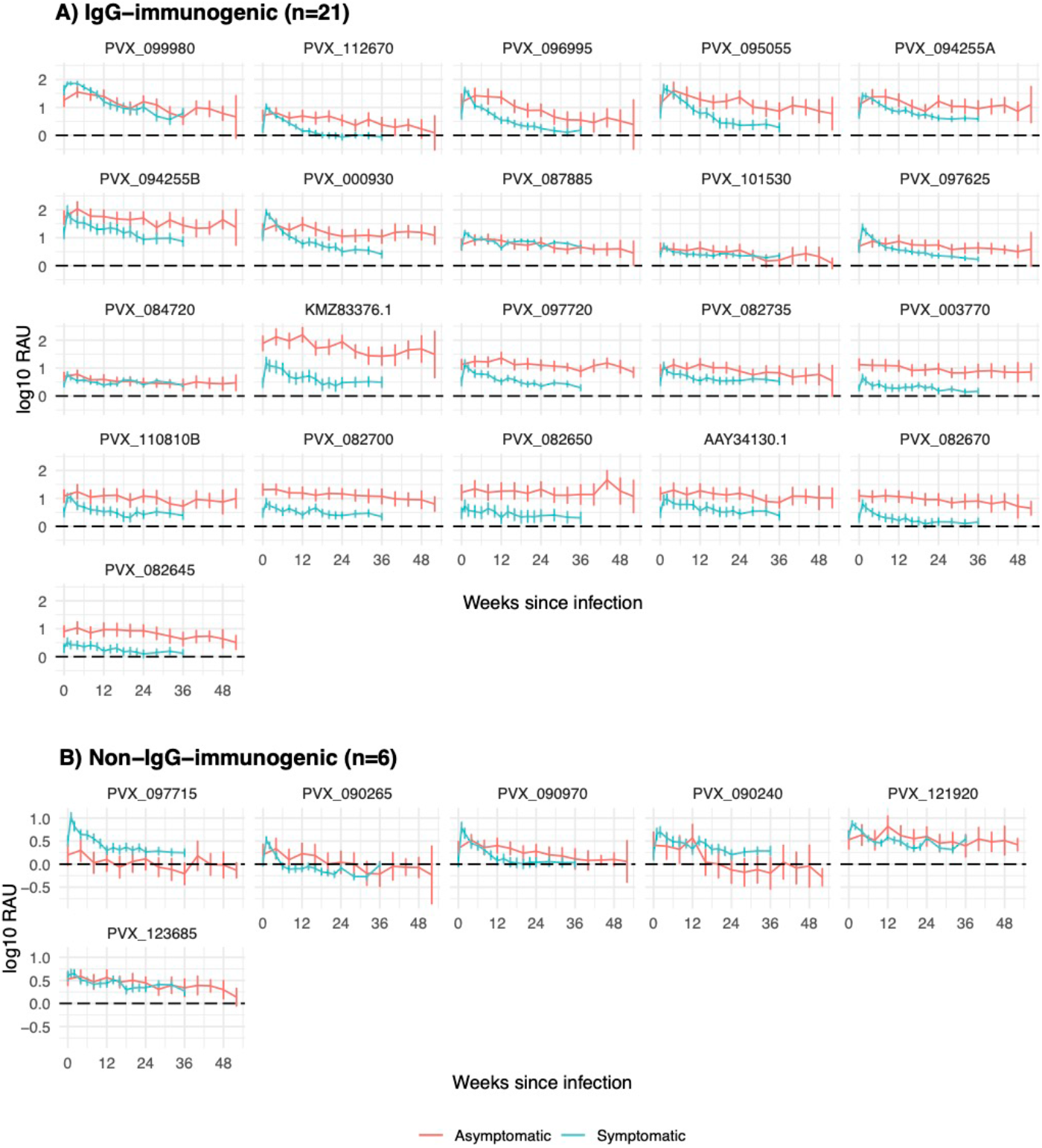
Comparison of adjusted total IgG against 27 *P. vivax* antigens following symptomatic or asymptomatic *P. vivax* infections. The kinetics of total IgG antibody against 27 IgG-immunogenic *P. vivax* antigens were determined following both asymptomatic (for a year) and symptomatic (for 9 months) *P. vivax* infections in Thai individuals in the absence of recurrent infections. Antibody data after asymptomatic infection were aligned to the time of infection, and thus sample size diminishes at later time points. The observed kinetic profiles were categorised into **A)** IgG-immunogenic (> 1 SD at 1-week post-infection above negative control baseline; n = 21) and **B)** non-IgG-immunogenic (> 1 SD at 1-week post-infection; n = 6) based on the asymptomatic profiles. Data were adjusted by subtracting the median of pooled negative control panels (n=274) and are expressed as median ± 95% CI.

**Figure S8.**
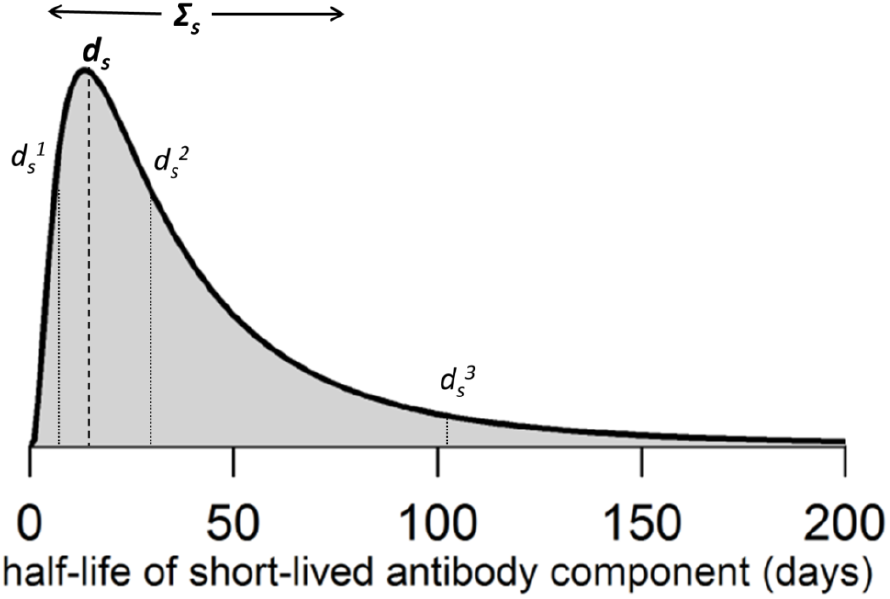
Schematic representation of the relationship between the population-level and individual-level parameters for the half-life of the short-lived ASCs. It is assumed that half-lives are log-Normally distributed throughout the population. The population-level parameters define the mean *d*_*s*_ and standard deviation Σ_s_ of this distribution. The individual-level parameters *d*_*s*_^*n*^ for each of the *n* participants follow the log-Normal distribution defined by the population-level parameters. Three representative individual-level parameters are shown for illustration.

